# Algal-derived extracts act as selective ecological filters shaping soil microbiomes, bacterial traits, and tomato performance under biotic stress

**DOI:** 10.64898/2026.03.30.710257

**Authors:** Millia Rose McQuade, Daniel Silva, Kusum Niraula, Ana Sofía Rodrigues dos Santos, Lucas Amoroso Lopes de Carvalho, Stela Jokic, Krunoslav Aladic, Ivana Flanjak, Igor Jerkovic, Inês Rebelo Romão, Joana do Carmo Gomes, Jelena Vladic, Juan Ignacio Vílchez

## Abstract

Modern agriculture faces the dual challenge of increasing food production while reducing reliance on synthetic inputs that degrade soil ecosystems and compromise long-term sustainability. Algal biomasses have emerged as promising biostimulants, yet their capacity to selectively modulate soil microbiomes and plant growth-promoting bacterial (PGPB) functions remains poorly understood. Here, we evaluated 17 phylogenetically and biochemically diverse macro- and microalgal extracts to determine their effects on soil microbial communities, bacterial functional traits, and tomato (*Solanum lycopersicum*) performance. Algal supplementation selectively restructured microbial communities without disrupting overall diversity, promoting taxa associated with plant-beneficial functions, including *Bacillus*, *Pseudomonas*, and Actinobacteria. In soil microcosms, specific treatments increased culturable bacterial abundance by up to ∼200-fold relative to the initial soil. Functional assays revealed strong extract- and strain-dependent responses. Siderophore production and ACC-associated activity were the most consistently stimulated traits, whereas auxin production, biofilm formation, and proline synthesis showed more variable or context-dependent responses. Notably, *Ulva* sp. (AP11.2) enhanced siderophore production across the majority of isolates, with over four-fold increases in individual strains, while *Arthrospira*-derived extracts (NG4.1, N14.1) consistently promoted bacterial growth across multiple taxa. In contrast, extracts such as *Nannochloropsis* sp. (NG6.1) and *Tetraselmis* sp. (NG5.1) induced more selective or inhibitory responses, highlighting extract-dependent functional trade-offs. Integration of biochemical and biological datasets identified fatty acid composition as a key axis associated with microbial functional responses, whereas volatile organic compound profiles showed weaker and less consistent associations. These microbiome and functional shifts translated into improved plant performance, with algal treatments increasing tomato growth and reducing mortality by approximately 20% under non-sterile soil conditions characterized by pathogen-associated pressure. Together, these findings demonstrate that algal extracts act as selective modulators of soil microbiomes, enhancing specific bacterial functions and improving plant performance in a context-dependent manner. This work provides a mechanistic framework for the development of targeted algal-based biostimulants aimed at reducing agrochemical inputs and advancing microbiome-informed agriculture.

## 1 INTRODUCTION

Soil degradation and the increasing demand for agricultural productivity are placing unprecedented pressure on global food systems. Current estimates indicate that nearly 40% of the global land surface is degraded, affecting over 3.2 billion people worldwide (Talukder et al., 2021; Vicente-Serrano et al., 2024). At the same time, agricultural production must increase by up to 70% by 2050 to meet rising food demand (Wijerathna-Yapa and Pathirana, 2022). Addressing this challenge requires a transition from input-intensive systems towards more sustainable approaches that harness biological processes rather than relying solely on synthetic inputs, which have contributed to soil degradation, biodiversity loss, and disruption of soil microbial communities (Ogidi and Akpan, 2023; D’Odorico et al., 2019). In this context, the soil microbiome has emerged as a central regulator of plant performance, influencing nutrient cycling, plant growth, and resilience to environmental stress. Rather than passive components of the soil environment, microbial communities actively shape plant outcomes through complex functional interactions, including phytohormone production, nutrient mobilization, and stress mitigation (Dellagnezze and Sierra-Garcia, 2023). Consequently, strategies that modulate microbiome composition and function represent a promising avenue for improving crop productivity in a sustainable manner.

Biostimulants have gained increasing attention as tools to enhance plant growth and resilience through biological mechanisms. These include a wide range of substances and microorganisms that act directly on plant physiology or indirectly through interactions with the soil microbiome (Yakhin et al., 2017; Buss et al., 2025). Among these, algal-derived biomasses are particularly promising due to their rich and diverse biochemical composition, including phytohormones, polysaccharides, and lipids. Macroalgae have a long history of agricultural use, while microalgae are increasingly recognized for their capacity to influence soil microbial diversity and function (Wang et al., 2021; Ferreira et al., 2024; Osorio-Reyes et al., 2023). Although algal extracts have been shown to enhance plant growth and modify microbial communities (Wang et al., 2018; Renaut et al., 2019), the mechanisms underlying these effects remain poorly understood. In particular, the links between algal biochemical composition, microbial community restructuring, and the modulation of specific plant growth-promoting bacterial (PGPB) traits are not well resolved. Most studies have examined plant responses or broad shifts in microbial composition in isolation, with limited integration of chemical, microbial, and plant-level data.

Addressing this gap requires a more integrated understanding of how algal-derived compounds influence microbial functional traits that directly impact plant performance. Plant growth-promoting bacteria contribute to nutrient acquisition, phytohormone production, and stress tolerance, yet their activity is highly context-dependent and influenced by environmental inputs. Identifying which biochemical features of algal biomasses drive these functional responses is therefore critical for the rational design of microbiome-based biostimulants. We hypothesized that algal extracts act as selective ecological filters that drive deterministic shifts in soil microbial communities, leading to trait-specific modulation of plant growth-promoting bacterial functions. Furthermore, we hypothesized that these effects are linked to specific biochemical features of algal biomasses, particularly fatty acid composition, which may influence microbial physiology and functional outcomes.

In this study, we evaluated 17 phylogenetically and biochemically diverse algal biomasses for their capacity to modulate soil microbiomes, influence plant growth-promoting bacterial functions, and enhance tomato (*Solanum lycopersicum*) performance. By integrating algal chemical profiling with microbial community analyses, functional trait assays, and in vivo plant experiments, this work provides a mechanistic framework for understanding algae-mediated microbiome modulation and supports the development of targeted, microbiome-oriented strategies for sustainable agriculture.

## 2 MATERIALS AND METHODS

### 2.1 Algal biomass selection and extract preparation

A total of sixteen dried and powdered algal biomass samples, comprising both microalgae and macroalgae, were supplied by the microalgal company Necton (Algarve, Portugal). In addition, one commercial biofertilizer product (Terralgae; Terraguar, Aveiro, Portugal), formulated from multiple algal species, was included in the study. The algal genus, corresponding sample codes, and application concentrations used in plant tests, soil microcosms, and bacterial assays are listed in Table 1.

**Table 1.**
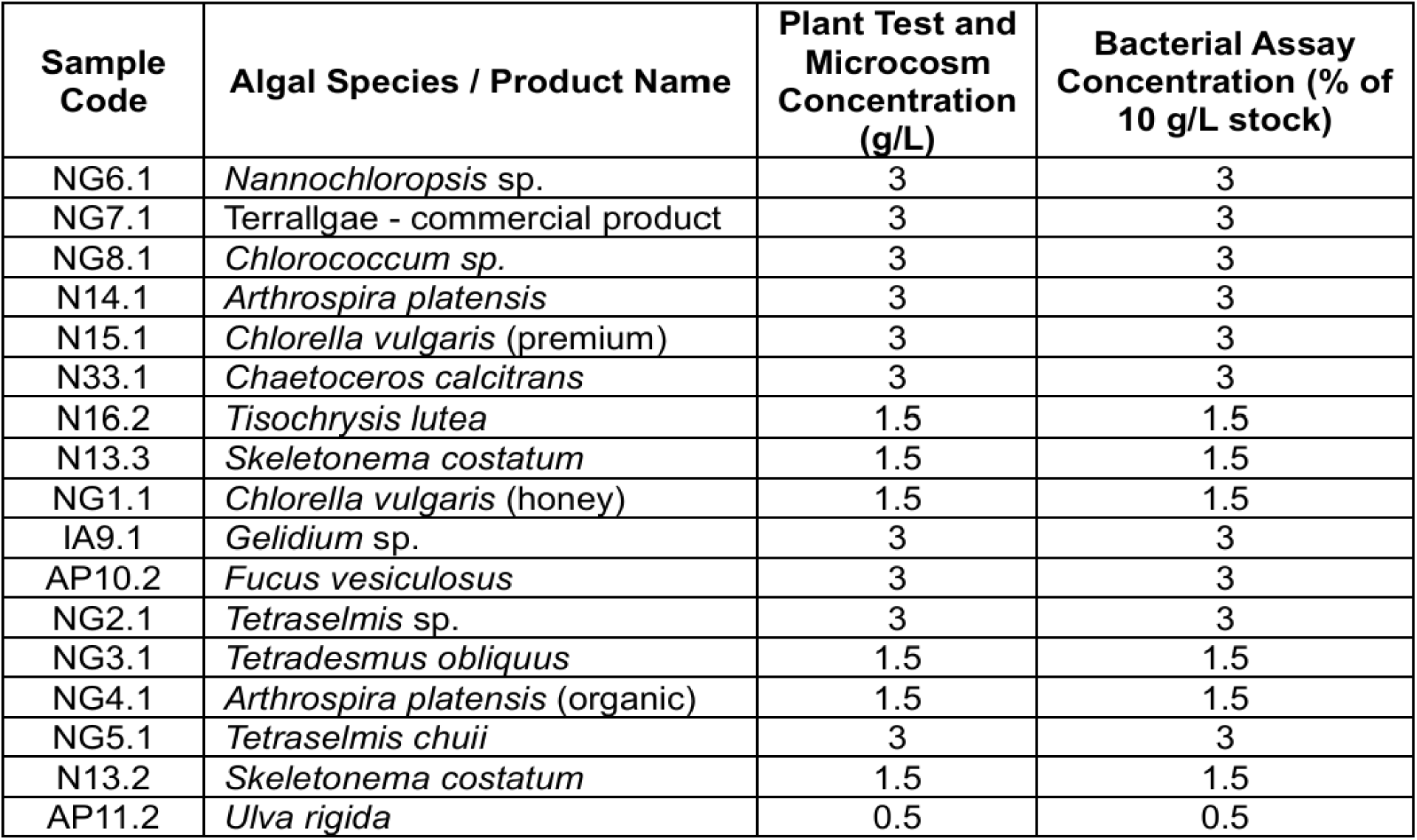
Algae biomass samples with their corresponding code names, as well as concentrations for plant tests and microcosms (g/L) and bacterial assays (% of 10 g/L stock).

For extract preparation, dried algal powders were suspended in double-distilled water (ddH₂O) at a concentration of 10 g/L. Suspensions were incubated under agitation (170 rpm) at 23 °C for 16–20 h to obtain aqueous extracts. Following extraction, suspensions were filtered through cloth to remove large particulate material and subsequently diluted with ddH₂O to the concentrations required for plant experiments, soil microcosms, and bacterial assays. These concentrations were selected based on previous optimization plant trials conducted by our collaborators. For bacterial assays requiring sterile conditions, extracts were filtered through 0.22 µm membrane filters prior to use.

### 2.2. Chemical characterization of algal biomasses

#### 2.2.1 Volatile organic compound analysis (HS-SPME/GC–MS)

Headspace volatile organic compounds (VOCs) were analyzed using solid-phase microextraction coupled to gas chromatography–mass spectrometry (HS-SPME/GC–MS). A manual SPME fibre coated with polydimethylsiloxane/divinylbenzene (PDMS/DVB, 65 µm; Supelco, Bellefonte, PA, USA) was used and conditioned according to the manufacturer’s instructions. For each analysis, 0.5 g of algal biomass was placed in a 20 mL glass vial, to which 2 mL of deionized water and 0.5 g NaCl were added to promote volatilization.

Vials were sealed with PTFE/silicone septa and incubated in a PAL 120 RSI autosampler equipped with a heating compartment. Samples were equilibrated at 40 °C for 15 min, followed by headspace extraction for 45 min. Following extraction, the SPME fibre was retracted, transferred to the GC injector, and thermally desorbed at 250 °C for 7 min. Analyses were performed using an Agilent 7890B gas chromatograph coupled to an Agilent 5977A mass spectrometer (Agilent Technologies, Palo Alto, CA, USA), equipped with an HP-5MS capillary column (30 m × 0.25 mm, 0.25 µm). The injector operated at 250 °C in split mode (1:5), with helium as the carrier gas at a constant flow rate of 1.0 mL min⁻¹. The oven temperature program started at 70 °C (2 min), ramped at 3 °C min⁻¹ to 200 °C, and was held at 200 °C for 18 min. The mass spectrometer operated in electron ionization mode (70 eV), scanning m/z 40–400. Injector and detector temperatures were set to 250 °C and 300 °C, respectively. Compound identification was based on comparison with Wiley 9 and NIST 17 mass spectral libraries, as well as retention indices calculated using C9–C25 n-alkanes. Each sample was analyzed in triplicate, and results are reported as mean values.

#### 2.2.2 Fatty acid analysis (GC–FID)

Total lipids were extracted using the Folch method with chloroform/methanol (2:1, v/v) (Folch et al., 1957). Preparation of fatty acid methyl esters (FAMEs) followed the methodology described in Jerković et al. (2021). FAMEs were analyzed using a Shimadzu GC-2010 Plus gas chromatograph equipped with a flame ionization detector (FID) and an SH-FAMEWAX™ capillary column (30 m × 0.32 mm i.d., 0.25 µm film thickness). The injector and detector temperatures were set to 240 °C and 250 °C, respectively. A 2 µL injection volume was used with a split ratio of 1:100. The oven temperature program was as follows: initial temperature 120 °C (held for 5 min), increased to 220 °C at 5 °C min⁻¹, and held for 20 min. Nitrogen was used as the carrier gas at a constant flow rate of 1.26 mL min⁻¹. Fatty acids were identified by comparison with a certified reference standard (Supelco F.A.M.E. Mix, C4–C24), analyzed under identical conditions. Results were expressed as the percentage of each fatty acid relative to total fatty acids.

### 2.3 Soil microcosm and culturable microbial analysis

Topsoil (0–15 cm) was collected from six tomato field sites in Vila Franca de Xira, Portugal (pathogen proliferation reported soil). Samples were sieved to remove debris and combined in equal volumes to produce a homogenized soil mixture representative of the initial microbial community. Baseline culturable bacterial abundance was estimated by suspending approximately 0.5 mL of mixed soil in 0.45% NaCl solution, followed by serial dilution (10⁻¹-10⁻⁴) and plating on LB agar. Plates were incubated at 28 °C for 24 h, after which colony-forming units (CFUs) were enumerated and normalized to dry soil weight. Morphologically distinct colonies were isolated and purified for further analysis. Microcosms were established by adding 4 mL of each algal extract (Table 1), water, or NPK fertilizer (3.825 mL/L) to 25 mL of homogenized soil in 50 mL conical tubes. Samples were thoroughly mixed and incubated for 7 days at room temperature.

Following incubation, culturable bacteria were re-isolated using the same serial dilution and plating approach. Changes in CFU abundance and relative composition were assessed relative to the initial soil. Morphologically distinct colonies exhibiting a relative abundance increase greater than 5% were selected for taxonomic identification. Genomic DNA was extracted using a heat-shock method (95 °C for 10 min), and the V5-V8 region of the bacterial 16S rRNA gene was amplified using primers 799F and 1392R. PCR products were verified by agarose gel electrophoresis and sequenced by Sanger sequencing (GENEWIZ, Germany). Taxonomic assignment was performed using BLAST against the NCBI database.

### 2.4 Metabarcoding analysis

#### 2.4.1 DNA extraction and sequencing

Metabarcoding analyses were performed on samples of the initial soil (T0) and soils collected from each microcosm after 7 days of incubation (T1). Samples were stored at −70 °C prior to DNA extraction. Total DNA was extracted using the DNeasy PowerSoil Pro Kit (Qiagen, Netherlands) according to the manufacturer’s instructions. DNA quantity and quality were assessed using a NanoDrop One/Oneᶜ microvolume spectrophotometer (Thermo Scientific, USA). Bacterial community composition was characterized by amplifying the V3-V4 region of the 16S rRNA gene using primers 341F and 806R. PCR amplification consisted of an initial denaturation at 95 °C for 3 min, followed by 25 cycles of denaturation (95 °C, 30 s), annealing (55 °C, 30 s), and extension (72 °C, 45 s), with a final extension at 72 °C for 10 min. Purified amplicons were pooled in equimolar concentrations and sequenced on an Illumina NovaSeq platform (2 × 250 bp paired-end) by Novogene (Cambridge, UK).

#### 2.4.2 Bioinformatic processing

Sequencing data were processed using QIIME2 (v2020.6). Quality filtering, denoising, chimera removal, and paired-end merging were performed using the DADA2 plugin. Based on quality profiles, forward and reverse reads were truncated at 280 bp and 210 bp, respectively. Amplicon sequence variants (ASVs) were inferred and retained for downstream analyses. Taxonomic classification was performed using a Naive Bayes classifier trained on the SILVA 138 reference database, trimmed to the V3-V4 region. Singleton and extremely low-abundance ASVs were removed prior to analysis.

#### 2.4.3 Community composition, diversity, and functional prediction

ASV tables were collapsed to class, family, genus, and species levels, and relative abundances were calculated per sample. Alpha diversity metrics (Chao1 richness, Shannon diversity, Simpson diversity, dominance, and Pielou’s evenness) were computed using QIIME2 and R (vegan package). Differences among treatments were assessed using ANOVA followed by Tukey’s HSD test. Beta diversity was evaluated using weighted UniFrac distances. Ordination analyses included PCoA, PCA, NMDS, and DCA, and hierarchical clustering was performed using UPGMA. Treatment effects were tested using PERMANOVA. Microbial functional potential was inferred using PICRUSt2, with predicted functions annotated against KEGG Orthology, KEGG pathways (Levels 1-3), enzyme commission numbers, COG, Pfam, and TIGRFAM databases. Community assembly processes were assessed using βNTI, phylogenetic normalized stochasticity ratio (pNST), and iCAMP.

### 2.5 Plant growth-promoting bacterial trait assays

Bacterial isolates were evaluated for plant growth-promoting traits including growth rate, biofilm formation, auxin production, ACC deaminase activity, proline production, and siderophore production using microplate-based assays. Strains were pre-cultured in LB broth (Miller’s modification, 20 g/L) at 28 °C under agitation (180 rpm) for 12–16 h. Cultures were standardized to an OD600 of 0.05 and distributed into 96-well plates with a final volume of 200 µL per well, consisting of bacterial culture, algal extract (Table X), and assay-specific components. Each assay included treatment, culture-only control, and extract-only control, and all conditions were performed in triplicate. Biofilm formation was quantified using a crystal violet assay (Coffey and Anderson, 2014). Auxin production was assessed using a modified Salkowski assay (Gang et al., 2019). ACC deaminase activity was inferred from growth in M9 medium supplemented with ACC as the sole carbon source (Naveed et al., 2008). Proline production was measured using a ninhydrin-based assay (Goswami et al., 2022), and siderophore production was quantified using the chrome azurol sulfonate (CAS) assay (Arora and Verma, 2017).

### 2.6 Tomato seedling assay

Three algal biomasses were selected for plant experiments: AP11.2 (*Ulva rigida*), N13.3 (*Skeletonema costatum*), and N16.2 (*Tisochrysis lutea*), based on previous screening trials and biochemical assay results. Extracts were applied at concentrations of 0.5, 1.5, and 1.5 g/L, respectively. Soil was collected and prepared as described above and mixed with turf at a ratio of 3:1 (v/v). To distinguish between direct and microbiome-mediated effects, half of the soil was tyndallized by heating at 70 °C for three consecutive days, with intermediate aeration periods to allow germination of resistant propagules. Tomato seeds (*Solanum lycopersicum*) were surface sterilized in 70% ethanol for 10 min, rinsed with ddH₂O, and soaked overnight. Seeds were sown directly into 0.25 L pots. Plants were watered daily with 40 mL of water. Algal extracts (40 mL) were applied three weeks after sowing, followed by a second application one week later. One week after the second application, plants were harvested, roots were washed, and root and shoot lengths were quantified using ImageJ (v1.54p). A two-way ANOVA followed by Tukey’s post hoc test was used to evaluate the effects of soil type, treatment, and their interaction.

### 2.7 Statistical and multivariate analyses

Volatile compound and fatty acid datasets were normalized to relative abundances (%) and log-transformed [log₁₀(x + 1×10⁻⁶)] to stabilize variance. Hierarchical clustering was performed using Euclidean distance and average linkage. Heatmaps were generated using the dendrogram order. Datasets of volatile compounds (VOCs) and fatty acids were arranged as feature-by-sample tables. Values were converted to per-sample relative abundances (%) summing to 100%, then log-transformed as log₁₀(x + 1×10⁻⁶) to stabilize variance and retain zeros. This scaling minimized dominance of high-abundance compounds and enabled comparison across samples. Samples were grouped by hierarchical clustering (Euclidean distance, average linkage), and heat maps were generated using the dendrogram order. For VOCs, class-specific panels and a compact map of the 30 most variable compounds (based on log-scale variance) were produced. All visualizations were created in Python (Matplotlib/SciPy). Warmer colors indicate higher relative abundance. Other statistical analyses were performed using GraphPad Prism (v10.6.0). Data are presented as mean ± standard deviation (n = 3), and differences among treatments were assessed using one-way ANOVA followed by Dunnett’s post hoc test (p < 0.05). Spearman’s rank correlations were calculated to assess relationships between biochemical profiles (VOCs and fatty acids) and bacterial functional traits. Correlations with |r| ≥ 0.5 were considered biologically relevant.

## 3. RESULTS AND DISSCUSION

### 3.1 Volatile organic compound profiles and sample clustering

Volatile organic compounds (VOCs) were characterized by HS-SPME/GC–MS, revealing a broadly conserved class-level composition across algal biomasses, dominated by terpenoid-derived compounds and ketones (Figure 1; Annex A1). Apocarotenoids and other oxygenated terpenoids accounted for approximately 10-43% of total identified volatiles and frequently represented the most abundant class. These included recurrent carotenoid cleavage products such as β-cyclocitral, safranal, α/β-ionone, 4-oxoisophorone, and dihydroactinidiolide, consistent with oxidative carotenoid metabolism in photosynthetic organisms (Simkin, 2021). Ketones constituted the second major class (8-33%), comprising both aliphatic enones (C7-C9) and cyclic ketones. Alcohols represented a smaller fraction (5-8%), with 1-octen-3-ol and hex-1-en-3-ol frequently detected. The presence of 1-octen-3-ol is consistent with linoleic acid oxidation and has been associated with antimicrobial activity in vitro (Xiong et al., 2017).

**Figure 1.**
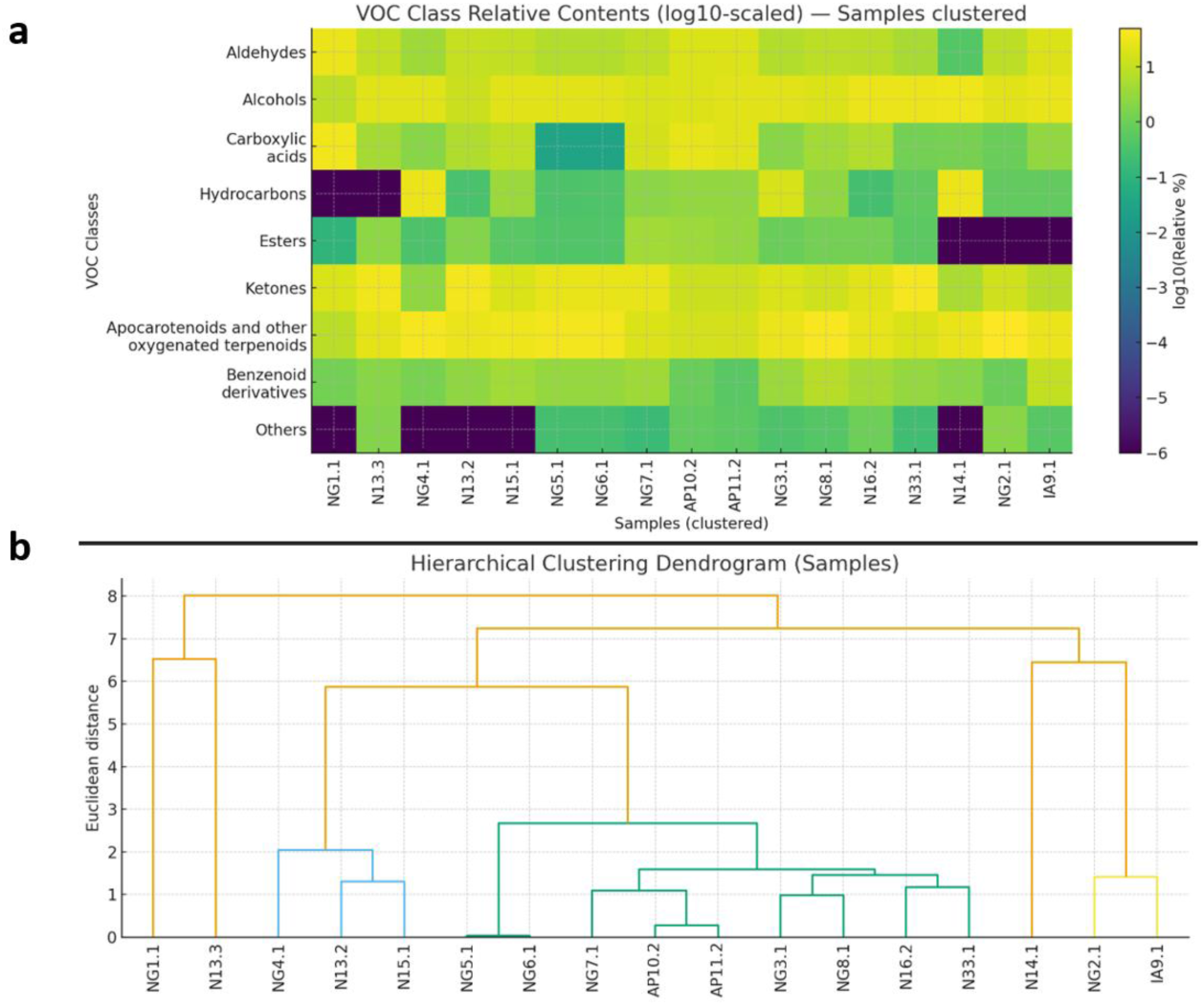
Heat-map clustering of VOCs class profiles. (a) Clustered heat-map displaying distribution of VOC classes across 17 algal samples. Values are log₁₀-transformed relative percentages and are shown on a color scale representing standardized relative abundance (b) Corresponding dendrogram indicating sample similarity based on Euclidean distance.

Aldehydes ranged from trace levels to 26% and were dominated by C6–C10 saturated and unsaturated species (e.g., hexanal, (E)-2-hexenal, nonanal, decanal), products of the LOX/HPL pathway involved in stress and defense signaling in photosynthetic organisms (Bate and Rothstein, 1998). Carboxylic acids were present at minor to moderate levels (0–30%), while hydrocarbons were generally low but reached 26–30% in a subset of samples. Esters and benzenoid derivatives were consistently minor components. Hierarchical clustering based on VOC class distributions revealed reproducible sample groupings (Figure 1). Several paired or closely related profiles were observed (e.g., AP10.2-AP11.2; NG5.1-NG6.1; NG3.1-NG8.1; N16.2-N33.1), indicating broadly similar volatile class patterns within these sets. In contrast, NG1.1 and N14.1 occupied distinct positions, reflecting divergent class distributions: NG1.1 was characterized by high acids and aldehydes with minimal hydrocarbons, whereas N14.1 displayed elevated hydrocarbons and alcohols but low aldehydes and esters. N13.3 was also distinctive, lacking hydrocarbons and showing a ketone-rich profile.

Reordering samples according to the dendrogram clarified broader compositional tendencies (Figure 1). A hydrocarbon-rich group emerged around NG4.1 and N14.1, while NG5.1 and NG6.1 clustered as low-acid samples enriched in ketones, alcohols, and apocarotenoids. A third group (AP10.2, AP11.2, NG7.1) was enriched in acids and aldehydes, whereas N16.2 and N33.1 exhibited alcohol- and ketone-rich profiles with moderate contributions from other classes. Despite these distinctions, most samples shared a conserved terpenoid–ketone backbone, limiting between-sample contrast at the VOC class level. Although many detected volatiles have been reported to exhibit biological activity, class-level VOC correlations with bacterial functional traits were weak and inconsistent. Apart from a recurring negative tendency between ketone abundance and biofilm formation in some comparisons, no robust or systematic associations were observed. This likely reflects both the compositional similarity across samples and the complexity of natural volatile mixtures, where individual compounds may exert opposing or context-dependent effects.

Taken together, these results indicate that VOC profiles are relatively conserved across algal biomasses at the class level, limiting their discriminatory power in explaining treatment-specific microbial and functional responses. This suggests that, under the conditions tested, VOC composition may play a secondary or context-dependent role compared to other biochemical fractions.

### 3.2 Fatty acid composition and compositional structuring

In contrast to VOCs, fatty acid (FA) profiles exhibited substantially greater compositional variability across algal biomasses (Figure 2; Annex A2). At the class level, saturated fatty acids (SFA) ranged from 24.4 to 61.3%, monounsaturated fatty acids (MUFA) from 6.6 to 48.9%, and polyunsaturated fatty acids (PUFA) showed the widest dispersion, with n-6 PUFA spanning 3.6-40.3% and n-3 PUFA 0.5-51.7%. Thus, the polyunsaturated fraction, particularly n-3 PUFA, represented the principal axis of between-sample variability.

**Figure 2.**
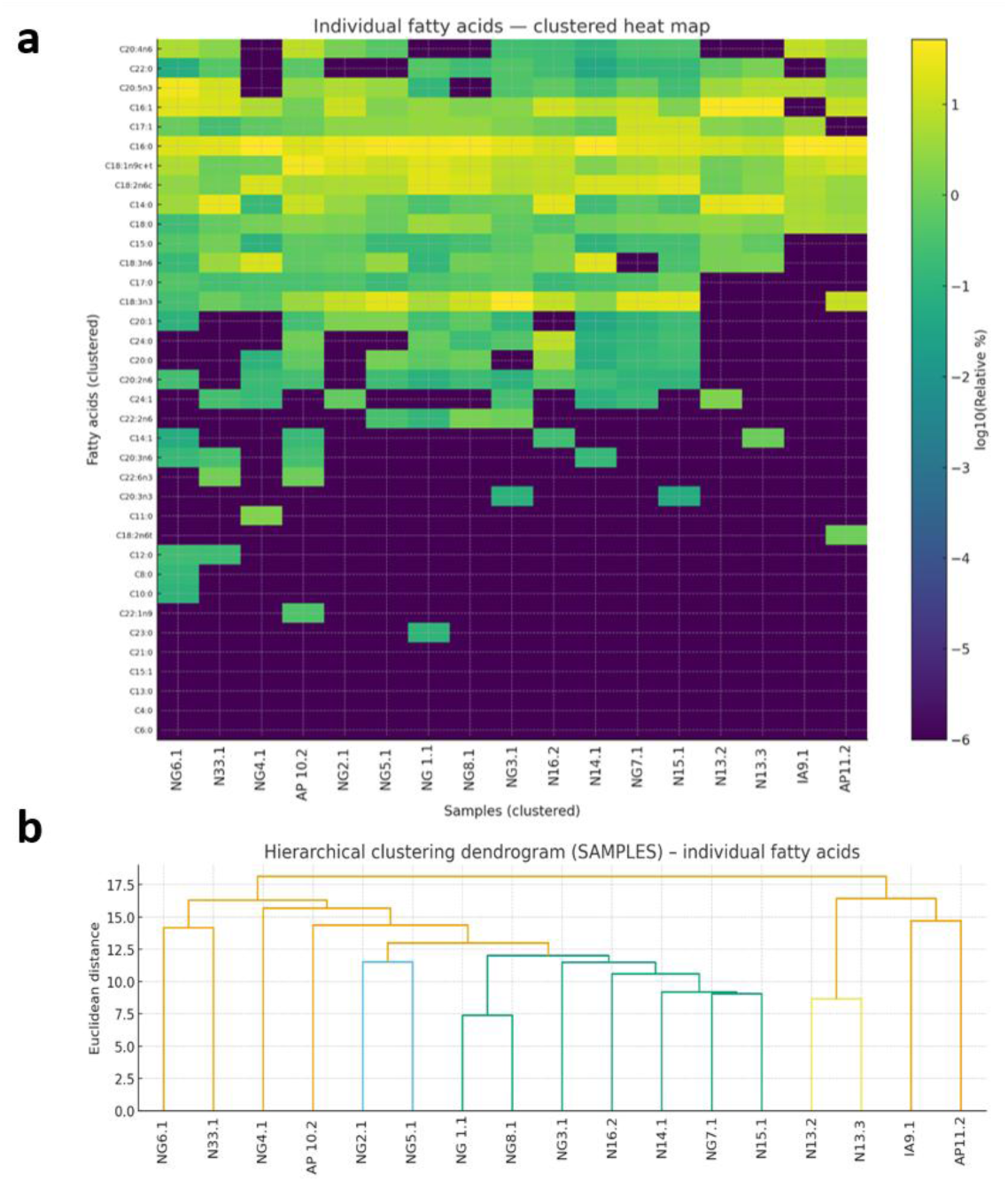
Heat-map clustering of fatty acid profiles. (a) Clustered heat-map displaying distribution of fatty acids across 17 algal samples. Values are log₁₀-transformed relative percentages and are shown on a color scale representing standardized relative abundance (b) Corresponding dendrogram indicating sample similarity based on Euclidean distance.

At the individual FA level, palmitic acid (C16:0) was consistently abundant and often the dominant component, ranging from approximately 15.5% to 50.0%. The principal MUFA were C16:1 and C18:1 n-9, with the latter varying from trace levels to 38.36%. Within PUFA, some samples were enriched in n-3 species, particularly α-linolenic acid (C18:3 n-3) and eicosapentaenoic acid (C20:5 n-3), whereas others were dominated by n-6 fatty acids such as linoleic acid (C18:2 n-6) and arachidonic acid (C20:4 n-6). Very-long-chain saturated fatty acids (C22:0-C24:0) were present at appreciable levels in only a limited subset of samples.

Hierarchical clustering based on the full FA matrix separated samples into three broad compositional tendencies (Figure 2): (i) n-3 PUFA-rich profiles with lower SFA; (ii) n-6-biased profiles with moderate SFA; and (iii) SFA/MUFA-dominated profiles characterized by elevated C16:1 and C18:1 n-9. This structuring provided a clearer and more discriminative chemical framework for interpreting downstream functional responses than VOC class distributions.

### 3.3 Taxonomic Responses to Algal Extracts

Metabarcoding analysis revealed clear microbial succession in the control microcosm over the incubation period (Figures 3 and 4). The initial soil community (T0) was dominated by Actinobacteria, with Bacilli and Proteobacteria present at lower abundances. Following incubation, the water control (T1) showed a marked shift towards increased relative abundance of Proteobacteria, while Bacilli declined significantly and Actinobacteria remained relatively stable. These trends are consistent with natural soil succession, where Proteobacteria and Actinobacteria outcompete Firmicutes (Fierer et al., 2007). The NPK treatment largely mirrored this trajectory, indicating that chemical fertilization reinforced existing community dynamics rather than inducing substantial compositional shifts. Although minor differences in specific taxa were observed, there was no clear enrichment of classical plant growth-promoting genera. These findings suggest that conventional fertilization maintains dominant microbial groups without substantially enhancing beneficial taxa, highlighting the need for alternative inputs such as algal biostimulants (Geisseler and Scow, 2014; Hartmann et al., 2014).

**Figure 3.**
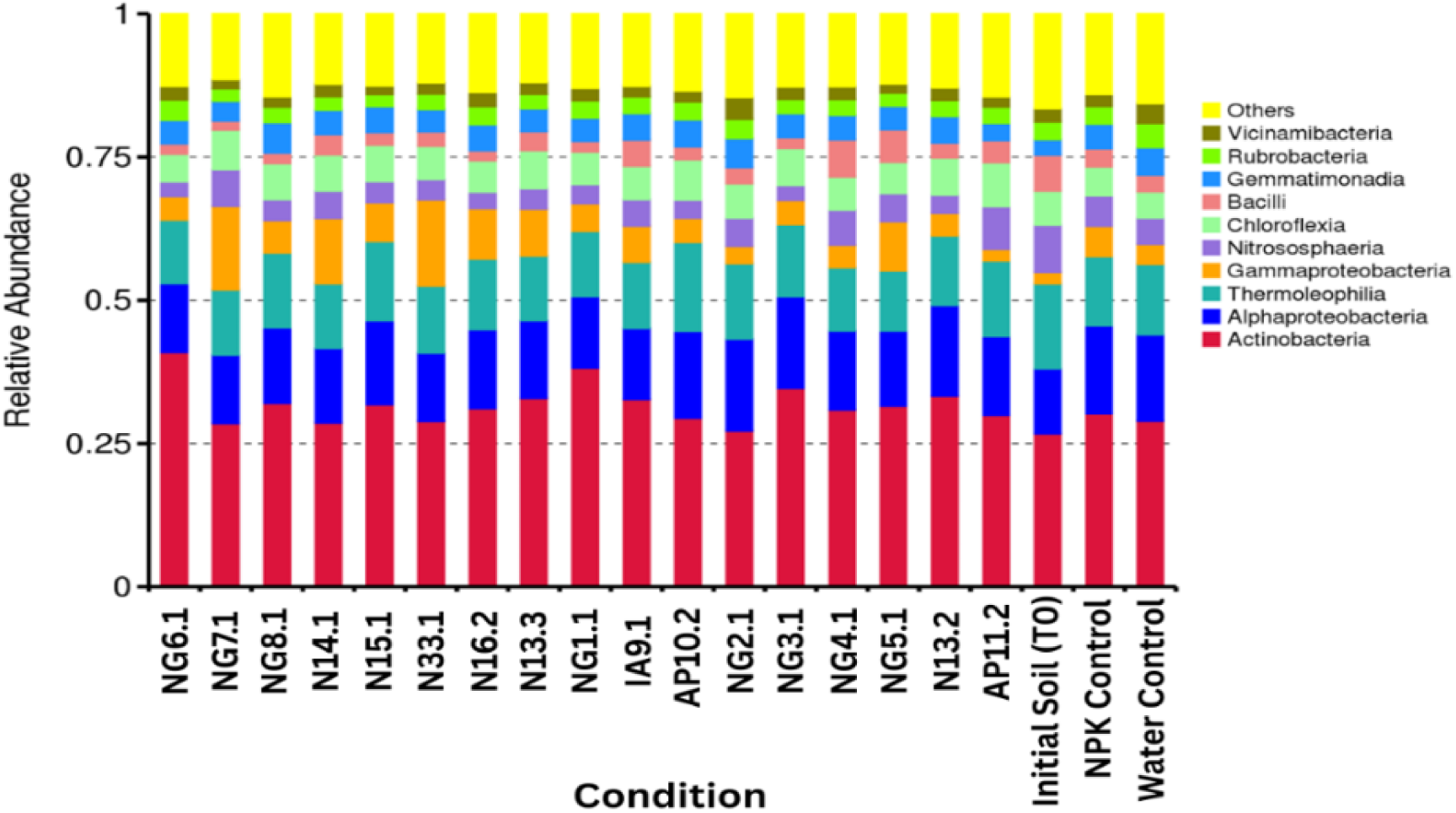
Relative abundance of the top 10 bacterial classes in algal microcosm treatments. Clustered stacked column chart displaying the relative abundances of the ten most dominant bacterial classes in algal supplemented and control microcosms (colour-coded as shown in the legend), along with a grouped category for all remaining classes (‘others’). ‘Initial soil (T0)’ represents the original soil community, while all other treatments, including algal applications and the water and fertilizer (NPK) controls, were assessed after 7 days (T1).

**Figure 4.**
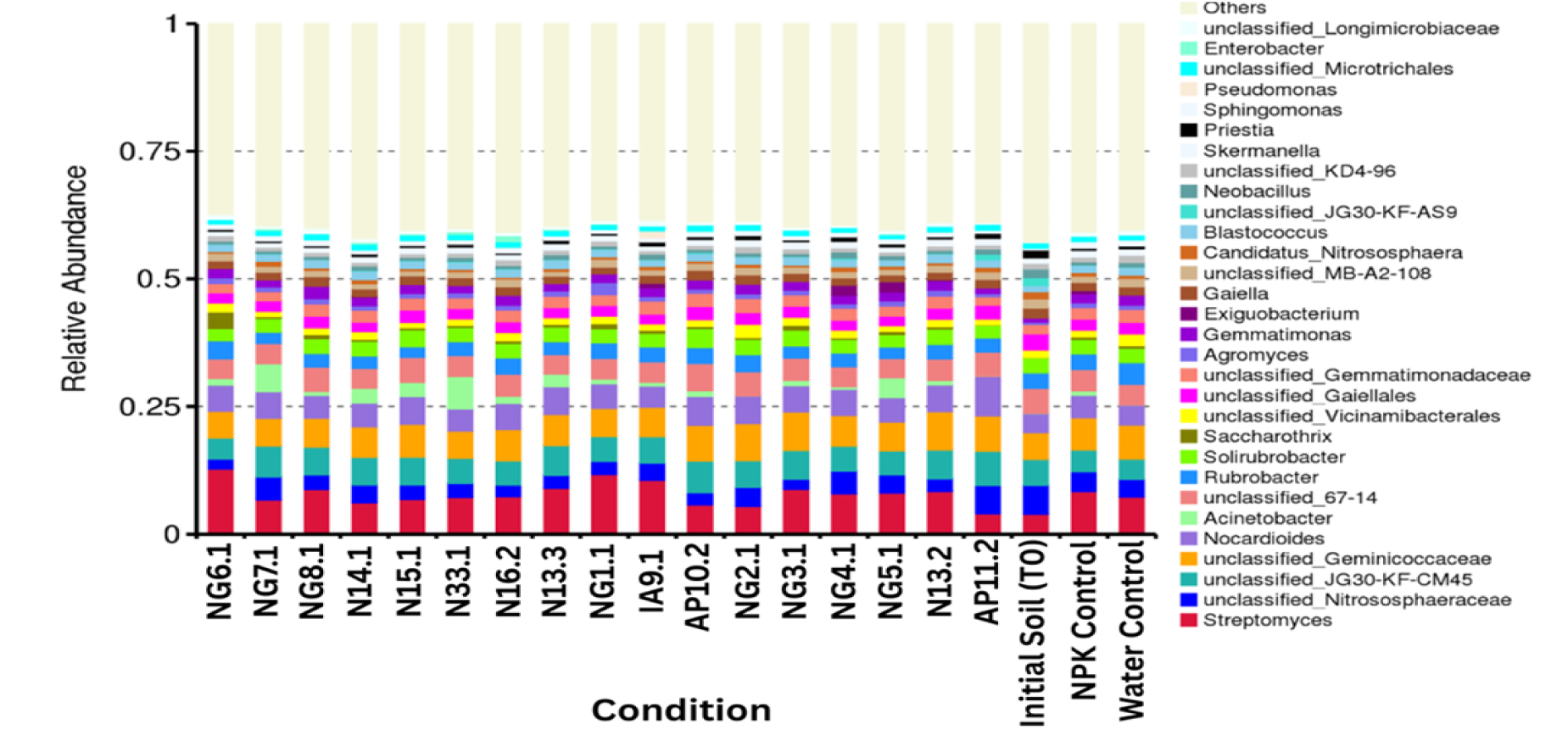
Relative abundance of the top 30 bacterial genera in algal microcosm treatments. Clustered stacked column chart showing the relative abundances of the 30 most abundant bacterial genera in algal supplemented and control microcosms, along with the combined abundance of the remaining genera grouped as ‘others’. ‘Initial soil (T0)’ represents the original soil community prior to treatment, while all other treatments, including algal applications and the water and fertilizer (NPK) controls, display the bacterial community after 7 days. The bacterial genera are color-coded as displayed in the legend.

In contrast, algal biostimulants induced distinct and treatment-specific shifts in community composition. For example, extract NG6.1 (*Nannochloropsis* sp.) markedly promoted Actinobacteria, increasing the relative abundance of key genera such as *Streptomyces*, Nocardioides, and *Saccharothrix*, taxa commonly associated with antibiotic production, pathogen suppression, and organic matter turnover (Chater et al., 2016; Ebrahimi-Zarandi et al., 2022). Conversely, N33.1 (*Chaetoceros* sp.) shifted the community towards a Proteobacteria-dominated state, with strong enrichment of *Acinetobacter* and the emergence of *Enterobacter*. N16.2 (*Tisochrysis* sp.) also promoted Enterobacter, suggesting enrichment of metabolically versatile taxa, although with potential implications for opportunistic behavior depending on environmental context (Bhattacharyya et al., 2017; Alzate Zuluaga et al., 2021). A distinct trajectory was observed for NG4.1 (*Arthrospira* sp.) and NG5.1 (*Tetraselmis* sp.), which restored Bacilli to levels comparable to the initial T0 soil. This shift was driven by increased abundances of *Neobacillus* and *Priestia*, genera frequently associated with plant growth promotion and stress resilience (Mmotla et al., 2025). Similarly, IA9.1 (*Gelidium* sp.) restored Bacilli while additionally promoting Pseudomonas, suggesting the potential establishment of synergistic consortia of key plant-beneficial taxa.

Notably, several of the enriched genera (e.g., *Bacillus*, *Pseudomonas*, and Actinobacteria) are well-documented producers of siderophores and other plant growth-promoting traits, providing a functional basis for the trait-level responses observed in subsequent assays. Collectively, these findings demonstrate that algal extracts can restructure soil microbial communities through selective enrichment of specific taxonomic groups, either reinforcing functionally relevant guilds or promoting alternative community configurations. While several treatments enhanced taxa linked to plant growth promotion, others favored less conventional groups, underscoring the importance of subsequent functional validation (e.g., biochemical assays) when evaluating their potential as biostimulants.

### 3.4 Alpha and Beta Diversity

At the community level, alpha diversity metrics indicated that algal supplementation generally preserved overall community structure (Figure 5). While several extracts were associated with reductions in Shannon diversity, such as NG6.1 (*Nannochloropsis* sp.), NG7.1 (Terrallgae), and N33.1 (*Chaetoceros* sp.), evenness remained consistently high across treatments, as reflected in stable Simpson’s indices. Conversely, extracts such as NG2.1 (*Tetraselmis* sp.) and AP10.2 (*Fucus* sp.) were associated with the highest richness and evenness, respectively, across treatments and controls, suggesting that certain algal extracts may enhance aspects of microbial diversity.

**Figure 5.**
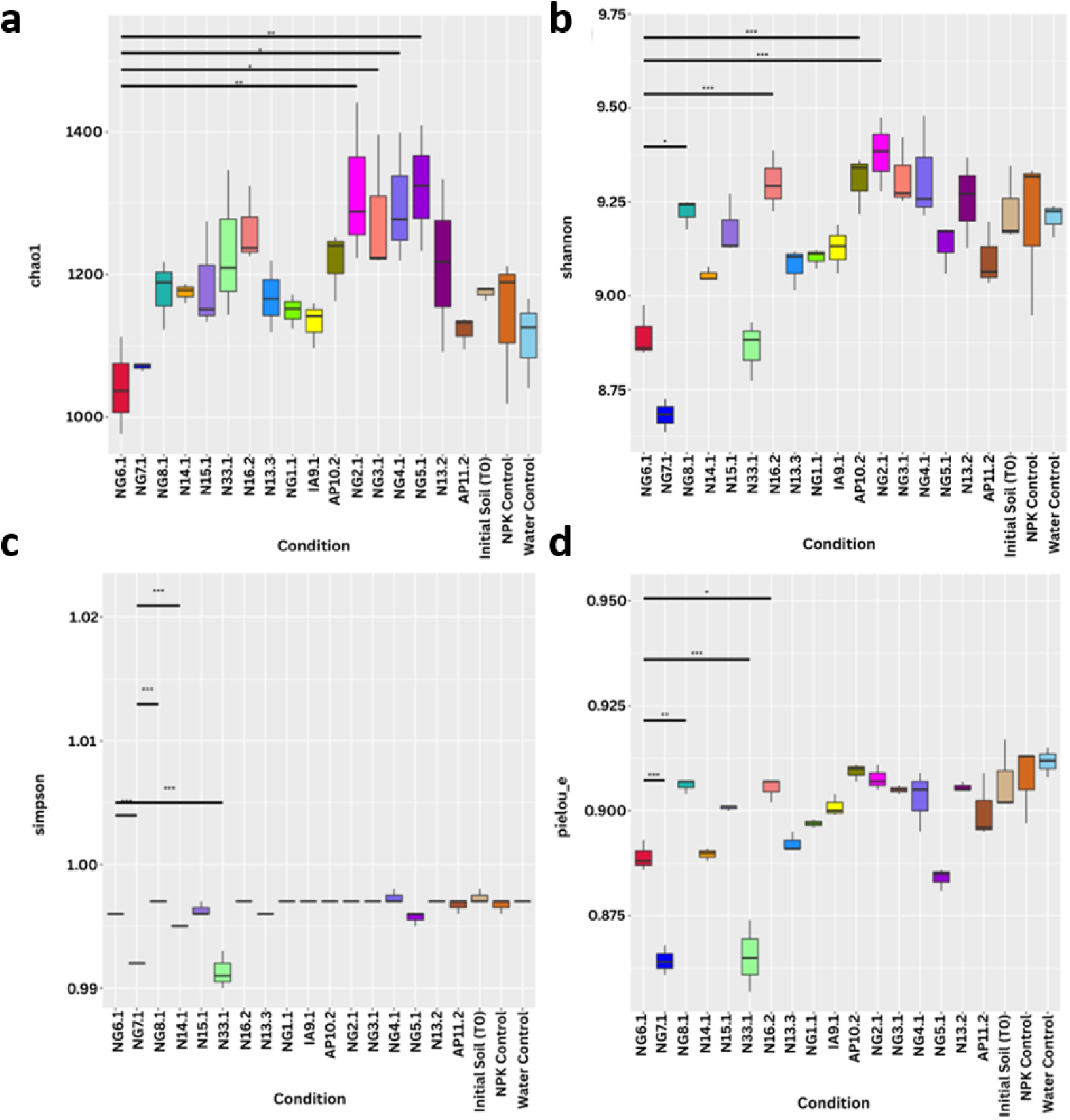
Alpha-diversity indices in algal microcosm treatments. Boxplots display the (a) Chao1, (b) Shannon diversity, (c) Simpson’s diversity, and (d) Pielou’s evenness indices of the initial soil community (Initial Soil [T0]) and of communities 7 days after application of aqueous algal extract or water and fertilizer (NPK) as controls. The lower, middle and upper lines of each box represent the first quartile (25th percentile), median (50th percentile), and third quartile (75th percentile), respectively, and the whiskers extend to the most extreme values within 1.5× the interquartile range from the first or third quartile. Statistically significant differences between treatments are indicated by asterisks (*p < 0.05, **p < 0.01, ***p < 0.001).

Beta diversity analyses further supported overall structural stability. Ordination approaches (including PCA, PCoA, NMDS, and DCA) showed that the majority of algal-treated communities clustered closely with the water control at T1, indicating that algal supplementation did not drive large-scale community divergence (Figure 6). However, NG6.1 (*Nannochloropsis* sp.) and IA9.1 (*Gelidium* sp.) exhibited greater separation in beta diversity space, consistent with their pronounced taxonomic enrichment patterns. Hence, these findings indicate that algal extracts can selectively restructure microbial communities while maintaining general community integrity. This combination of targeted compositional shifts without widespread disruption is a key property of effective microbial biostimulants.

**Figure 6.**
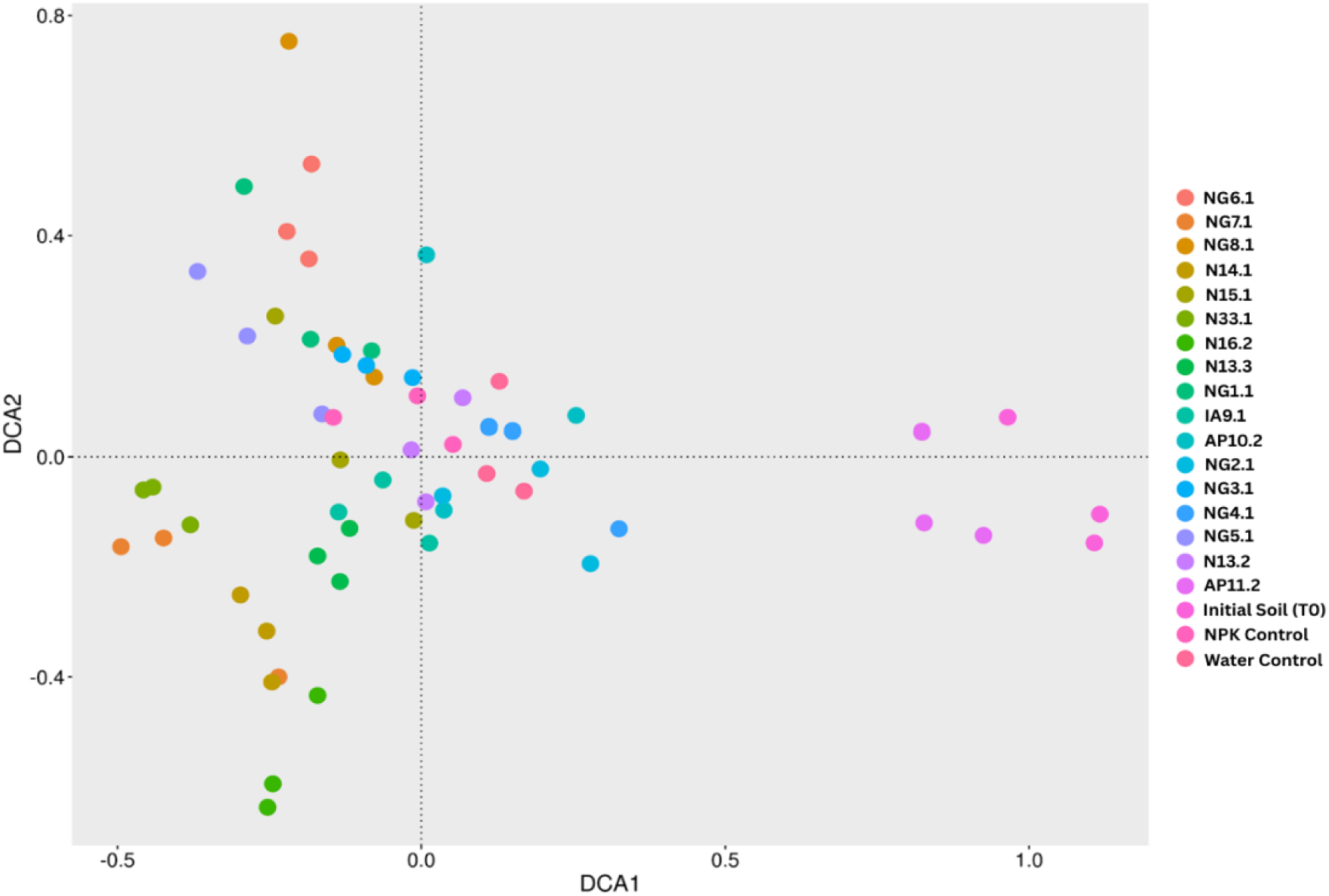
Beta-diversity DCA analysis of microcosm treatments. Ordination plot from Detrended Correspondence Analysis (DCA) showing differences in bacterial community composition among 17 algal supplemented microcosms and water and NPK (fertilizer) controls. Each microcosm includes three replicates (same-colored circles), with closer points representing more similar communities. DCA1 and DCA2, expressed as standard deviation units, correspond to the first and second largest gradients of beta diversity.

### 3.5 Functional Potential Shifts

Functional predictions using PICRUSt2 provided insight into the potential functional consequences of the observed community restructuring in response to algal supplementation. At the broadest level (KEGG level 1), metabolism remained the dominant functional category across all treatments (>80%), indicating that algal supplementation did not disrupt core microbial functional potential. At finer resolution, extract-specific functional enrichments were observed. For example, N33.1 (*Chaetoceros* sp.) increased pathways related to lipid metabolism, xenobiotic degradation, and membrane transport, consistent with the enrichment of copiotrophic taxa such as *Acinetobacter* and *Enterobacter*.

In contrast, NG6.1 (*Nannochloropsis* sp.) enhanced functions associated with secondary metabolite biosynthesis, including ansamycin production, reflecting the increased abundance of Actinobacteria such as *Streptomyces* and *Saccharothrix*. Similarly, NG4.1 and IA9.1 (*Arthrospira* sp.) were associated with enrichment of amino acid metabolism and tricarboxylic acid (TCA) cycle pathways, consistent with the metabolic versatility of Bacilli and *Pseudomonas* enriched under these treatments. Importantly, these functional profiles are inferred from taxonomic composition and therefore represent predicted functional potential rather than direct measurements of microbial activity. As such, while the observed patterns are consistent with the ecological roles of the enriched taxa, they should be interpreted as indicative rather than definitive evidence of functional shifts.

Within this context, the alignment between taxonomic enrichment and predicted functional categories suggests that algal extracts modulate microbiome functional potential in ecologically coherent ways. The observed enrichments are consistent with processes relevant to nutrient cycling, stress tolerance, and pathogen suppression. Comparable functional responses have been reported for other organic amendments and biostimulants (Backer et al., 2018; Rouphael and Colla, 2020). However, the treatment-specific and guild-associated patterns observed here highlight the capacity of algal extracts to act as targeted modulators of microbiome functional potential.

### 3.6 Community Assembly Processes

Null-model analyses revealed that community assembly in algal extract treatments was predominantly governed by deterministic processes (Figure 7). βNTI values were consistently strongly negative, indicating homogeneous selection as the dominant driver of community structuring. In agreement, pNST values remained below the stochasticity threshold of 0.5, further supporting a predominance of deterministic assembly. iCAMP analysis provided a more detailed partitioning of assembly processes, attributing most community turnover to homogeneous selection (47.6% ± 3.0), with secondary contributions from drift (37.8% ± 3.3) and dispersal limitation (∼12.3%). Homogenizing dispersal contributed minimally (≤3%), while heterogeneous selection was negligible.

**Figure 7.**
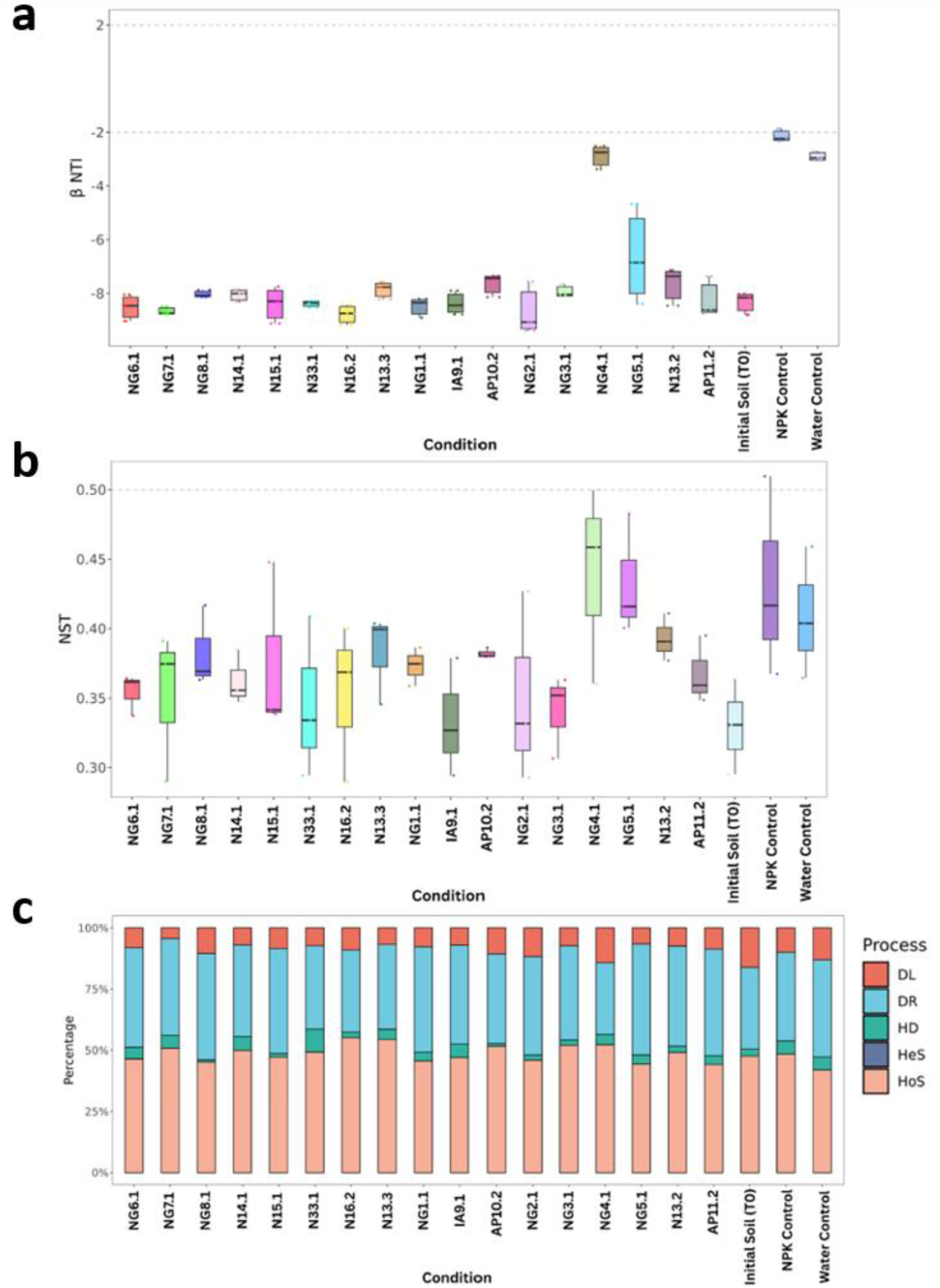
Community Assembly Analysis of Microcosm treatments. βNTI, pNST, and iCAMP analyses across algal extract, water, and NPK microcosm treatments. (a) Boxplots of β-nearest taxon index (βNTI) for pairwise comparisons between replicates within each treatment. Values < -2 indicate homogeneous selection, while values between -2 and +2 indicate stochastic assembly. (b) Boxplot of phylogenetic Normalized Stochasticity Ratio (pNST) for pairwise comparisons between replicates within each treatment. Values < 0.5 indicate predominantly deterministic assembly, while values > 0.5 indicate predominantly stochastic assembly. (c) Percent stacked column chart of iCAMP analysis showing the relative contribution of five ecological processes (homogeneous selection [HoS], heterogeneous selection [HeS], homogenizing dispersal [HD], dispersal limitation [DL], and drift [DR]) to community assembly within each treatment.

Together, these results indicate that algal extracts reinforce the existing soil environmental filter, promoting selective enrichment rather than stochastic community restructuring. The dominance of homogeneous selection suggests that algal-derived inputs impose consistent ecological pressures that favor specific, functionally coherent microbial guilds, such as Actinobacteria or Bacilli. In contrast to untreated or NPK-amended soils, which typically exhibit higher stochasticity, algal-treated systems displayed more constrained and reproducible assembly trajectories. This increased determinism may be advantageous in agronomic contexts, as it implies more predictable enrichment of beneficial microbial groups.

### 3.7 Population Assay

From the composite soil mix at T0 and the microcosms after 7 days of incubation (T1), a total of 54 morphologically distinct culturable isolates were recovered through plating and morphological characterization. Relative abundance profiles across treatments revealed clear shifts in culturable community composition following algal supplementation (Figure 8). Based on an abundance threshold of >5% increase in algal treatments relative to controls, 16 isolates were selected for taxonomic identification via 16S rRNA gene sequencing (Table 2). Importantly, the taxonomic identities of the isolated strains broadly aligned with the dominant or enriched taxa identified through metabarcoding, supporting the representativeness of the culturable fraction for downstream functional characterization.

**Figure 8.**
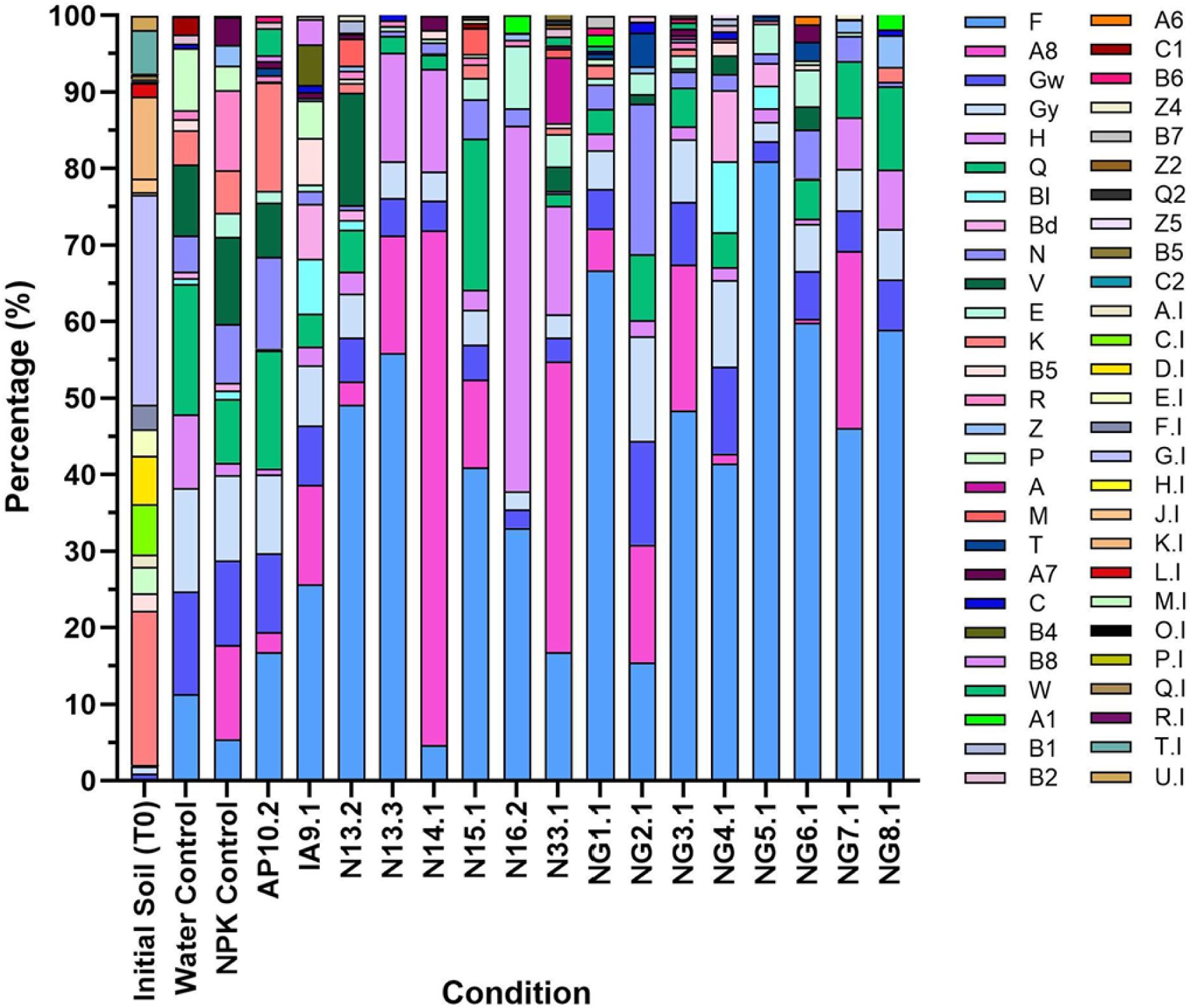
Population distribution of culturable morphologically distinct isolates. Stacked column chart showing the relative abundances of morphologically distinct culturable isolates (codes indicated in the legend) from original tomato field soil and in algal-treated microcosms, including water and NPK (universal fertilizer) control microcosms after seven days.

**Table 2.** Identification of morphologically distinct isolates from microcosm analysis. Sixteen selected morphologically distinct isolates (each with an assigned code name) were identified to species level based on 16S rRNA gene sequences using the BLAST database. The sequence similarity values between each isolate and the closest database match are also reported.

| Strain | Identification | BLAST<br>Threshold (%) |
| --- | --- | --- |
| A8 | *Bacillus sanguinis* | 99.85 |
| B4 | *Exiguobacterium acetylicum* | 98.96 |
| BD | *Exiguobacterium acetylicum* | 99.71 |
| E | *Peribacillus frigoritolerans* | 98.69 |
| F | *Enterobacter quasihormaechei* | 98.85 |
| GW | *Priestia aryabhattai* | 98.97 |
| H | *Leclercia adecarboxylata* | 99.00 |
| K | *Enterobacter quasihormaechei* | 99.70 |
| T | *Bacillus australimaris* | 99.71 |
| V | not identified |  |
| Z | *Priestia megaterium* | 99.11 |
| P | *Priestia aryabhattai* | 99.56 |
| BI | *Exiguobacterium acetylicum* | 99.13 |
| A | not identified |  |
| Q | not identified |  |
| GY | *Peribacillus frigoritolerans* | 99.85 |

All isolates were assigned to genera with well-documented plant growth-promoting bacteria (PGPB), including *Bacillus*, *Priestia*, *Peribacillus*, *Leclercia*, *Exiguobacterium*, and *Enterobacter*, which are known to exhibit traits such as phytohormone production, nutrient solubilization, and siderophore synthesis (Delegan et al., 2021; Fanai et al., 2024; Mmotla et al., 2025). Although several isolates exhibited very high 16S rRNA sequence similarity, distinct colony morphologies justified their retention as separate isolates for downstream functional assays, pending whole-genome sequencing.

Among all isolates, strain F (*Enterobacter quasihormaechei*) was the most dominant, being detected across all treatments and reaching a peak relative abundance of 80.98% in the NG5.1 (*Tetraselmis*) treatment. This dominance coincided with NG5.1 exhibiting the lowest Shannon and Simpson diversity indices, reflecting the strong prevalence of *E. quasihormaechei*. More broadly, alpha diversity analysis indicated that control treatments (initial soil, water, and NPK) maintained higher culturable diversity than most algal-treated microcosms, suggesting that algal supplementation imposes selective pressures on culturable bacterial communities.

Notably, NG5.1 also yielded the highest abundance of culturable bacteria, averaging 25,924.33 ± 5,029.80 CFU/mg, in contrast to the initial soil mixture (T0), which exhibited the lowest abundance at 129.81 ± 40.50 CFU/mg (Figure 9). This indicates that NG5.1 supported substantial proliferation of *E. quasihormaechei*, both in relative proportion and absolute abundance. This pronounced dominance suggests that algal-derived compounds may create selective niches that favor specific bacterial taxa, leading to strong restructuring of the culturable soil microbiome.

**Figure 9.**
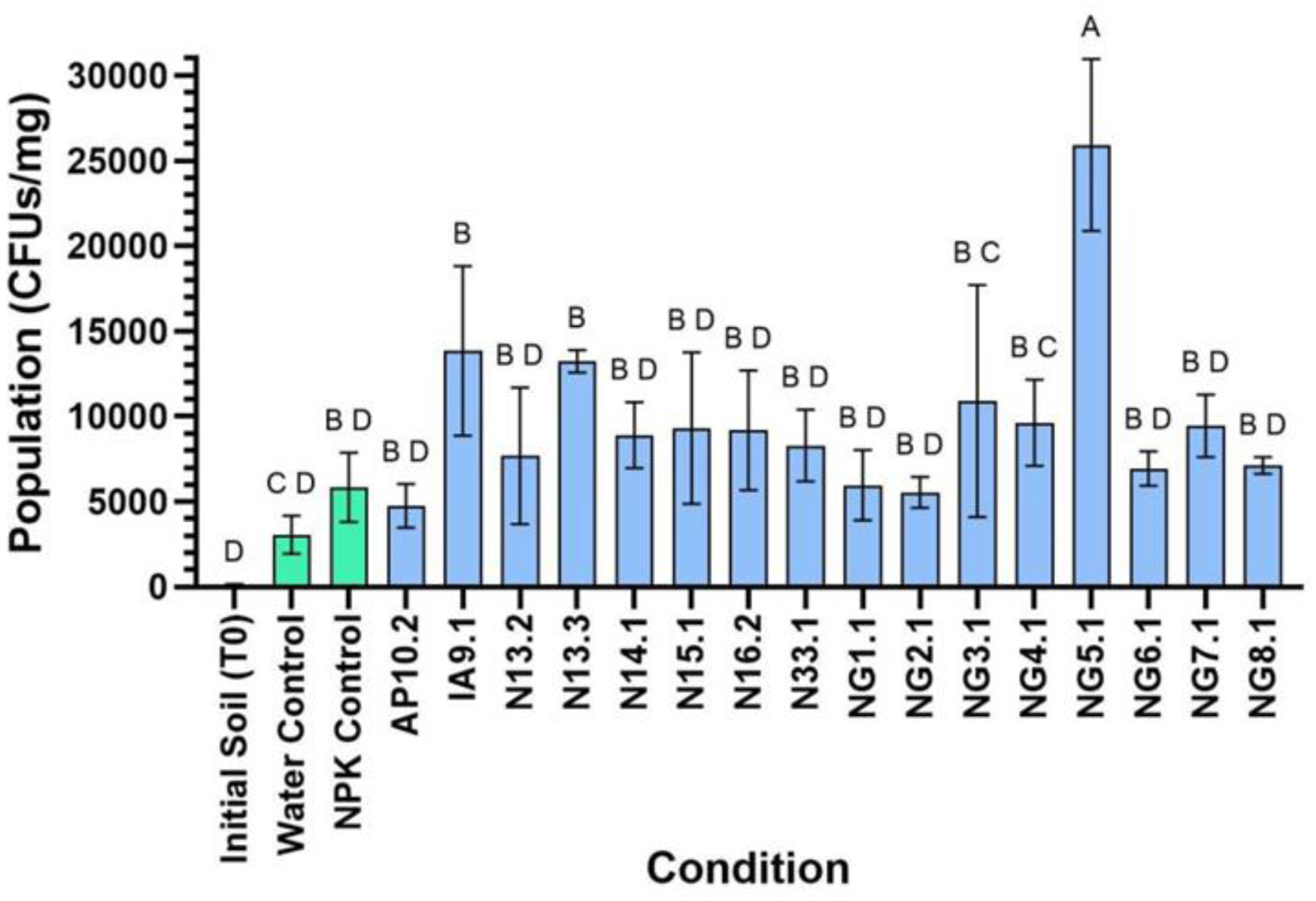
Abundance of culturable bacteria across microcosm treatments. Column graph displaying the average (n = 3) colony-forming units (CFUs) per milligram of soil dry weight for algal-supplemented, water, and NPK (fertiliser) microcosms. Error bars represent ±1 standard deviation. Data were analysed using one-way ANOVA followed by Dunnett’s post hoc test, with distinct letters indicating statistically significant differences (p < 0.05).

This culturomics-based approach enables the isolation of representative strains for functional characterization of plant growth-promoting traits, complementing metabarcoding analyses that capture total community diversity. Integrating these approaches provides a framework for linking community-level shifts with experimentally validated functional traits, facilitating the identification of candidate PGPB for future agricultural applications.

### 3.8 Effect Over Isolates Growth

Bacterial growth in response to aqueous algal extracts was assessed over 24 h by measuring optical density at 600 nm (OD₆₀₀) (Supplementary Figure 1). All isolates demonstrated the capacity to grow in the presence of algal treatments, although responses varied substantially depending on both the extract and the bacterial strain. Extracts NG4.1 and N14.1, both derived from *Arthrospira* sp., consistently promoted higher bacterial growth compared to other algal treatments and the non-supplemented control across the majority of isolates. This stimulatory effect is consistent with previous reports describing *Arthrospira*-derived biomasses as biostimulatory, likely due to their nutrient-rich composition and diverse bioactive compounds (Begum et al., 2024).

In contrast, several algal extracts exhibited inhibitory effects on bacterial proliferation, particularly NG6.1 (*Nannochloropsis* sp.), NG5.1 (*Tetraselmis* sp.), and NG7.1 (Terrallgae commercial mix). For example, isolate T (*Bacillus australimaris*) reached an OD₆₀₀ of approximately 1.5 under NG4.1 but plateaued at ∼0.5 under NG5.1, illustrating pronounced extract-dependent variation in growth responses. These inhibitory effects may reflect the presence of bioactive or antimicrobial metabolites, including long-chain polyunsaturated fatty acids, which have been reported to exert bacteriostatic or bactericidal effects depending on concentration and microbial sensitivity (Zourro et al., 2024; Carrillo and Anchundia, 2024). Importantly, such effects are likely to be highly context- and strain-dependent, and may contribute to selective pressures shaping microbial community composition rather than representing generalized toxicity. Hence, these findings indicate that algal extracts can differentially modulate bacterial growth, promoting or constraining proliferation in a strain- and extract-specific manner. This selective modulation is consistent with the compositional shifts observed in microcosm experiments, supporting a role for algal-derived compounds in shaping microbial community structure through differential growth responses.

### 3.9 Biofilm production

Biofilm formation, a key trait for bacterial surface attachment and root colonization, exhibited strong intrinsic variability among isolates (Ajijah et al., 2023). In addition to this inherent variability, algal supplementation significantly influenced biofilm formation across all 16 isolates (Supplementary Figure 2), with responses varying in both direction and magnitude depending on the specific extract–strain combination. In general, inhibitory effects predominated, with 104 significant reductions compared to 30 enhancements, indicating that most algal treatments reduced biofilm formation relative to controls. However, these responses were highly context-dependent and varied substantially across isolates and extracts. N14.1 (*Arthrospira* sp.) enhanced biofilm formation in the greatest number of isolates, consistent with its stimulatory effects on bacterial growth. This may reflect its nutrient-rich composition supporting the synthesis of extracellular polymeric substances required for biofilm development (Abdel Latif et al., 2022).

In contrast, N13.2 (*Skeletonema* sp.) exhibited the strongest inhibitory effects, reducing biofilm formation in 9 of the 16 isolates, suggesting the presence of bioactive compounds that interfere with biofilm development. Importantly, the predominance of inhibitory responses does not necessarily indicate a detrimental effect on plant-associated functions. Biofilm formation is a complex and tightly regulated process, and its reduction under specific conditions may reflect shifts in bacterial lifestyle strategies (e.g., planktonic versus surface-associated growth) or competitive dynamics among strains. As such, modulation of biofilm formation by algal extracts is likely to contribute to selective microbial structuring rather than representing a uniform suppression of functional potential. Collectively, these findings demonstrate that algal supplementation differentially modulates bacterial biofilm formation in a strain- and extract-dependent manner, with potential implications for rhizosphere colonization and plant-microbe interactions.

### 3.10 Auxins production

Auxin (indole-3-acetic acid, IAA) is a key phytohormone involved in root development and plant growth promotion (Pappalettere et al., 2024). Its production varied significantly among isolates, both intrinsically and in response to algal supplementation. However, compared to other traits, significant responses were less frequent and of lower magnitude, with only 13 increases and 51 decreases observed across all treatment–isolate combinations (Supplementary Figure 3). Algal supplementation induced both stimulatory and inhibitory responses within the same treatment. For example, NG6.1 (Nannochloropsis sp.) elicited both increases and decreases in IAA production depending on the isolate, highlighting strong strain-specific effects. In contrast, some extracts, such as NG1.1 (Chlorella sp.), had minimal overall impact despite showing more pronounced effects on other traits such as biofilm formation.

Notably, no consistent relationship was observed between biofilm formation and auxin production, indicating that these traits are likely regulated independently under the experimental conditions tested. Overall, the relatively limited responsiveness of IAA biosynthesis to algal supplementation suggests that auxin production may be more tightly regulated than other plant growth-promoting traits, and therefore less sensitive to modulation by algal-derived metabolites under the conditions tested.

### 3.11 Siderophores production

Siderophores are iron-chelating compounds that enhance the acquisition of bioavailable iron and support plant growth under iron-limiting conditions (Li et al., 2023). In contrast to other plant growth-promoting traits, siderophore production was consistently stimulated by algal supplementation across the majority of isolates, with 58 instances of significant increases compared to 27 decreases (Supplementary Figure 4).

Among the treatments, AP11.2 (*Ulva* sp.) showed the strongest overall stimulatory effect, significantly increasing siderophore production in 11 of the 16 isolates. Isolate A8 (*Bacillus sanguinis*) exhibited the greatest magnitude of response under this treatment, with more than a four-fold increase relative to the control. Across all isolates, strain T (*Bacillus australimaris*) displayed the highest responsiveness, with significant increases observed under all algal treatments (p < 0.001). The consistency and frequency of stimulatory responses distinguish siderophore production from other evaluated traits, which showed more variable or predominantly inhibitory patterns. This suggests that algal-derived compounds may broadly promote bacterial iron acquisition mechanisms, either by acting as metabolic substrates or by modulating regulatory pathways associated with iron limitation.

However, while the observed responses are robust, the underlying mechanisms remain unresolved and may involve indirect effects on cellular metabolism or iron availability rather than direct induction of siderophore biosynthesis pathways. Thus, these results indicate that siderophore production represents one of the most consistently enhanced bacterial functions in response to algal supplementation, highlighting it as a key functional axis through which algal extracts may influence plant–microbe interactions.

### 3.12 Proline production

Proline is an osmoprotectant associated with the stabilization of cellular structures under abiotic stress conditions such as drought and salinity (Zulfiqar and Ashraf, 2023). As with other plant growth-promoting traits, proline production was significantly influenced by algal supplementation. Overall, inhibitory effects predominated, with 149 instances of significant reduction compared to 58 increases across all isolate–treatment combinations (Supplementary Figure 5).

The strongest inhibitory effects were observed for treatments NG5.1 (*Tetraselmis* sp.), NG6.1 (*Nannochloropsis* sp.), and NG7.1 (Terrallgae), which also corresponded to the most pronounced reductions in bacterial growth observed in growth curve analyses. This convergence suggests that reduced proline production may, at least in part, reflect broader constraints on bacterial metabolism under these treatments. In contrast, AP11.2 (*Ulva* sp.) and N16.2 (*Tisochrysis* sp.) demonstrated more consistent stimulatory effects across multiple isolates, enhancing proline production. These patterns align with their positive effects on siderophore production and bacterial growth, suggesting coordinated modulation of multiple plant growth-promoting traits under specific algal treatments. Importantly, reduced proline production does not necessarily imply diminished plant-beneficial potential, as proline synthesis is typically associated with stress responses and may be downregulated under more favorable growth conditions. Therefore, the observed variation in proline production likely reflects shifts in bacterial physiological state rather than uniform suppression of functional capacity. These findings indicate that algal extracts differentially modulate proline production in a context-dependent manner, with effects linked to broader metabolic responses and growth dynamics.

### 3.13 Semi-Quantitative Determination of ACC Deaminase

ACC deaminase reduces plant ethylene levels by degrading 1-aminocyclopropane-1-carboxylate (ACC), thereby supporting continued root development and nutrient uptake under stress conditions (Hausinger et al., 2023; Gamalero et al., 2023). Bacterial ACC deaminase activity, inferred from growth using ACC as the sole carbon source, varied markedly across isolates in response to algal supplementation, although the overall trend was towards stimulation (Supplementary Figure 6).

Growth was significantly enhanced in approximately 50% of the isolate–treatment combinations (135/272), compared to only 15 significant reductions. NG4.1 (*Arthrospira* sp.) showed the strongest stimulatory effect, enhancing growth in 13 of the 16 isolates, whereas AP11.2 (*Ulva* sp.) exhibited the highest number of inhibitory responses (five). All isolates displayed basal growth on ACC in the absence of algal supplementation, indicating the presence of ACC deaminase activity, although growth levels remained low relative to those observed in nutrient-rich media.

Importantly, as this assay infers ACC deaminase activity from bacterial growth, the presence of additional carbon sources in algal extracts may confound interpretation. Enhanced growth may therefore reflect increased metabolic activity supported by exogenous substrates rather than direct upregulation of ACC deaminase activity. Consequently, while the results indicate strong isolate- and treatment-dependent responses, they should be interpreted as indicative of potential modulation of ACC-related metabolism rather than definitive evidence of changes in ACC deaminase enzymatic activity. In general, these findings suggest that algal extracts can influence bacterial growth under ACC-utilizing conditions, although direct enzymatic measurements would be required to confirm specific effects on ACC deaminase activity. Collectively, these functional assays indicate that algal extracts modulate bacterial activity in a trait-specific and context-dependent manner, rather than uniformly enhancing plant growth-promoting functions.

### 3.14 Associations between fatty acids and bacterial functional traits

Spearman correlation analyses revealed selective and target-dependent associations between fatty acid (FA) composition and bacterial functional readouts, rather than uniform relationships across traits (Annex Tables). Among the most consistent patterns, higher proportions of C15:0 were repeatedly associated with reduced biofilm formation, whereas biomasses enriched in C20:2 n-6 or containing moderate levels of C16:0 tended to support increased biofilm production. These patterns are consistent with a membrane-centric interpretation, whereby shorter or odd-chain saturated fatty acids increase membrane rigidity and may hinder early surface adhesion, while longer or partially unsaturated acyl chains facilitate membrane remodeling and energy supply via β-oxidation, conditions favorable for biofilm development. Such interpretations align with established links between acyl-chain composition, membrane fluidity, and biofilm physiology (Dubois-Brissonnet et al., 2016).

For ACC-related responses, positive associations were more frequently observed with n-6 C18–C20 fatty acids, whereas short-chain saturated fatty acids (C14:0–C15:0) tended to correlate with reduced activity. This pattern is consistent with evidence that ACC deaminase-associated metabolism is sensitive to cellular energy and redox balance, including oxygen-dependent regulation in plant-associated bacteria (Singh et al., 2015). Siderophore production showed recurrent positive associations with C18:1 n-9 and, in some cases, C24:0, suggesting that MUFA-rich or neutral-lipid-rich matrices may better support the energetic demands of siderophore biosynthesis. In contrast, auxin and proline responses were highly isolate-dependent, consistent with the notion that exogenous fatty acids can act as nutrients, signaling molecules, or stressors depending on uptake capacity, metabolic routing, and membrane homeostasis.

No single fatty acid emerged as a universal predictor of bacterial functional responses. Instead, the data define compositional “windows” associated with specific functional outcomes: biomasses enriched in C15:0 may be less favorable for biofilm formation or ACC-related responses; MUFA-rich or moderately n-6-enriched profiles (e.g., C18:1 n-9, C20:2 n-6) appear more supportive of siderophore production and ACC-associated activity; and very-long-chain saturated fatty acids (C22:0–C24:0) may exert context-dependent effects, particularly for auxin and proline responses. These associations support a mechanistic framework in which fatty acid composition influences bacterial physiology through membrane properties, metabolic routing, and energy availability, thereby shaping functional responses.

### 3.15 Integration with treatment-level biological responses

When considered alongside treatment-level biological outcomes, the compositional patterns identified in fatty acid (FA) profiles were broadly consistent with observed microbial and functional responses. The most growth-promoting extracts, NG4.1 and N14.1 (both *Arthrospira platensis*), exhibited FA profiles aligned with those associated with enhanced bacterial growth and ACC-related responses, including moderate C16:0 and/or enrichment in n-6 C18–C20 fatty acids such as C20:2 n-6. In contrast, extracts associated with reduced bacterial growth (NG5.1, NG6.1, NG7.1) corresponded to FA profiles predicted to increase membrane rigidity (e.g., short- or odd-chain saturated fatty acids) or to impose metabolic constraints, potentially through increased susceptibility to lipid peroxidation in the case of specific n-6 PUFA. These patterns are consistent with the inhibitory effects observed in both growth and proline assays, suggesting that FA composition contributes to shaping bacterial physiological responses under these treatments. Similarly, the strong and consistent stimulation of siderophore production observed for AP11.2 (*Ulva rigida*) aligned with its enrichment in C18:1 n-9 and, in some cases, C24:0, supporting the association between MUFA-rich profiles and enhanced siderophore biosynthesis. Conversely, certain n-6 PUFA (e.g., C20:3 n-6) were associated with reduced siderophore output, further highlighting the specificity of FA–function relationships. Importantly, these associations should be interpreted as indicative rather than causal, as multiple biochemical components may contribute to the observed biological effects, and interactions between compounds are likely to influence microbial responses. In particular, while FA composition provided a clearer axis for interpreting functional outcomes, other metabolite classes such as VOCs may exert context-dependent or combinatorial effects that are not resolved at the class level.

Across traits, responses were consistently strain- and extract-dependent, with no uniform enhancement across all functional readouts. For example, while siderophore production was broadly stimulated, other traits such as auxin production and biofilm formation showed variable or predominantly inhibitory responses. These patterns indicate that algal extracts do not act as generalised stimulants of microbial activity, but rather as selective modulators of specific functional traits and microbial subpopulations. This selective modulation is consistent with the deterministic community assembly patterns observed in microcosm experiments, where algal supplementation promoted reproducible enrichment of specific microbial guilds rather than stochastic community restructuring. Together, these results support a model in which algal extracts act as ecological filters, shaping both microbial composition and functional potential through differential growth responses and metabolic modulation.

Finally, it is important to note that several functional responses were inferred from proxy measurements (e.g., growth-based assays or predictive metagenomics), and therefore represent approximations of microbial activity under the tested conditions. While the overall coherence between chemical composition, microbial shifts, and functional assays strengthens the interpretation, direct mechanistic validation would be required to fully resolve causal relationships. This convergence of chemical composition, microbial community restructuring, and functional responses supports a predictive framework in which algal biomasses can be rationally selected or designed to target specific microbiome functions.

### 3.16 Plant Test

To evaluate the effects of algal supplementation on tomato growth, three selected extracts - AP11.2 (*Ulva* sp.), N13.3 (*Skeletonema* sp.), and N16.2 (*Tisochrysis* sp.) - were applied under optimized growth conditions in both untreated (original) and heat-sterilized (tyndallized) soils. Clear differences in plant survival were observed between soil types. In original soil, high mortality (50–70%) occurred across treatments, with symptoms consistent with soilborne pathogen infection (Chinheya et al., 2024; Cui et al., 2021; Zhang et al., 2023). The isolation of fungal taxa including *Fusarium longifundum*, *Rhizopus arrhizus*, and *Mucor circinelloides* from symptomatic plants further support that pathogen pressure contributed to the observed effects. In contrast, all plants grown in tyndallized soil survived, confirming effective pathogen suppression through sterilization. Despite pathogen pressure, algal treatments significantly improved plant performance. In original soil, all extracts increased root and shoot length relative to controls, with N13.3 and N16.2 also reducing mortality by ∼20% (figure X). In tyndallized soil, algal supplementation primarily enhanced shoot length and, for N13.3 and N16.2, significantly increased fresh weight (Supplementary Figure 7 and Supplementary 1).

These findings support a dual mode of action for algal extracts: a direct nutritional effect, evident under reduced microbial conditions, and an indirect, microbially mediated or stress-alleviating effect under biotic stress evident in the original soil (Chanthini et al., 2023). Overall, algal supplementation enhanced tomato growth, with the greatest relative benefits observed under biotic stress. This highlights the importance of plant-microbe interactions in mediating algal biostimulant effects and supports their potential as sustainable tools for improving crop resilience. The stronger relative effects observed under non-sterile conditions, together with the microbial and functional shifts described above, are consistent with a microbiome-mediated component contributing to plant performance, although direct causal relationships cannot be fully resolved within this study.

## 4 Conclusions

Modern agriculture’s reliance on synthetic inputs is driving environmental degradation and threatening long-term sustainability, highlighting the need for alternative strategies such as algal-derived biostimulants. In this context, the present study demonstrates that algal biomasses can enhance plant performance through both direct and microbiome-mediated mechanisms. Across 17 phylogenetically and biochemically diverse extracts, algal supplementation selectively restructured soil microbial communities while preserving overall community stability. These shifts were largely governed by deterministic assembly processes, indicating that algal inputs act as consistent ecological filters that promote the enrichment of specific microbial guilds.

At the treatment level, clear differences in biological responses were observed. *Arthrospira*-derived extracts (NG4.1 and N14.1) consistently promoted bacterial growth and supported multiple functional traits, while *Ulva* sp. (AP11.2) exhibited the strongest and most consistent stimulation of siderophore production across isolates. In contrast, extracts such as NG5.1 (*Tetraselmis* sp.), NG6.1 (*Nannochloropsis* sp.), and NG7.1 showed more selective or inhibitory effects, highlighting the importance of extract composition in determining microbial outcomes. At the functional level, algal extracts exerted trait-specific and isolate-dependent effects. Among the evaluated traits, siderophore production and ACC-associated responses showed the most consistent stimulation, whereas other functions such as auxin production, biofilm formation, and proline synthesis displayed more variable or context-dependent responses. These findings indicate that algal extracts do not act as generalized stimulants of microbial activity, but rather as selective modulators of specific bacterial functions. Fatty acid composition emerged as a key biochemical axis associated with microbial functional responses, providing a mechanistic link between algal biomass composition and microbiome modulation. In contrast, VOC profiles showed more conserved patterns and weaker associations with functional outcomes, suggesting a more limited role at the class level under the conditions tested.

Importantly, plant assays confirmed that algal supplementation improves tomato growth and survival, with the strongest relative effects observed under conditions of high biotic stress. In particular, treatments N13.3 (*Skeletonema* sp.) and N16.2 (*Tisochrysis* sp.) reduced plant mortality and enhanced growth under non-sterile conditions, supporting a microbiome-mediated or stress-alleviating effect. Under reduced microbial pressure, algal treatments primarily enhanced growth-related parameters, consistent with direct plant responses. While functional predictions and growth-based assays provide valuable insight into microbiome responses, they represent indirect measures of microbial activity. Therefore, the functional interpretations presented here should be considered indicative, and further mechanistic validation will be required to resolve causal relationships.

In summary, this work establishes algal extracts as targeted modulators of soil microbiomes, capable of enhancing beneficial bacterial functions and improving plant performance. These findings support their potential as sustainable tools for reducing reliance on synthetic inputs and advancing microbiome-informed agricultural practices. Importantly, the variability in responses across extracts highlights the need for targeted selection of algal biomasses based on desired microbiome and functional outcomes.

## Author Contributions

Conceptualization, Jelena Vladic and Juan Ignacio Vilchez; methodology, Millia Rose McQuade, Stela Jokic, Jelena Vladic and Juan Ignacio Vilchez; software, Millia Rose McQuade, Lucas Amoroso Lopes de Carvalho, Jelena Vladic and Juan Ignacio Vilchez; validation, Millia Rose McQuade, Daniel Silva, André Sousa, Inês Romão, Joana do Carmo Gomes, Stela Jokic and Kusum Niraula; formal analysis, Millia Rose McQuade, Jelena Vladic and Juan Ignacio Vilchez; investigation, Daniel Silva, André Sousa, Inês Romão, Millia McQuade, Kusum Niraula and Juan Ignacio Vilchez; resources, Jelena Vladic and Juan Ignacio Vilchez; data curation, Juan Ignacio Vilchez; writing—original draft preparation, Millia Rose McQuade, Jelena Vladic and Juan Ignacio Vilchez; writing—review and editing, Millia McQuade, Kusum Niraula, Jelena Vladic and Juan Ignacio Vilchez; visualization, Jelena Vladic and Juan Ignacio Vilchez; supervision, Jelena Vladic and Juan Ignacio Vilchez; project administration, Jelena Vladic and Juan Ignacio Vilchez; funding acquisition, Jelena Vladic and Juan Ignacio Vilchez. All authors have read and agreed to the published version of the manuscript.

## Acknowledgements

The authors thank ITQB NOVA (NOVA University of Lisbon, Oeiras, Portugal), the Faculdade de Ciências e Tecnologia of Universidade NOVA, and the GREEN-IT Research Unit for access to greenhouse facilities and supporting infrastructure. The authors also acknowledge Necton for providing the algal material, Hubel Verde for supplying the tomato farm soil used in this study, and Dr. Margarida Oliveira (GREEN-IT/ITQB NOVA) for her support and guidance.

---

**Supplementary Figure 1.**
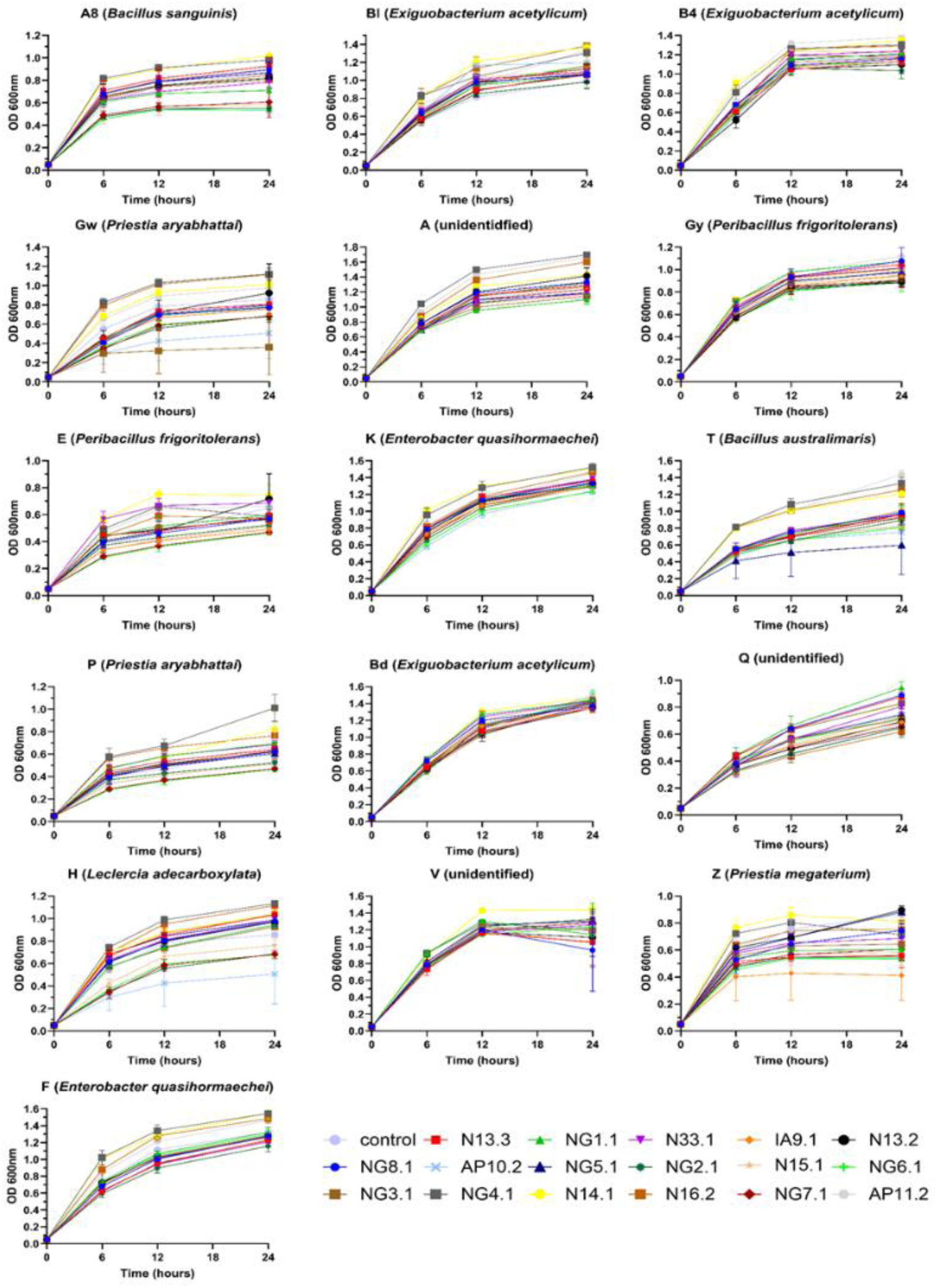
Growth analysis of bacterial isolates. Line graphs display the average (n = 3) growth patterns of 16 bacterial isolates in LB medium over 24 hours, with and without algal supplementation, inferred from optical density at 600 nm (OD₆₀₀). Each isolate is displayed in a separate graph, with treatment conditions indicated by different colours and symbols (as shown in the legend). OD₆₀₀ was measured at 0, 6, 12, and 24 hours, with the initial OD₆₀₀ = 0.05. Error bars represent ± one standard deviation.

**Supplementary Figure 2.**
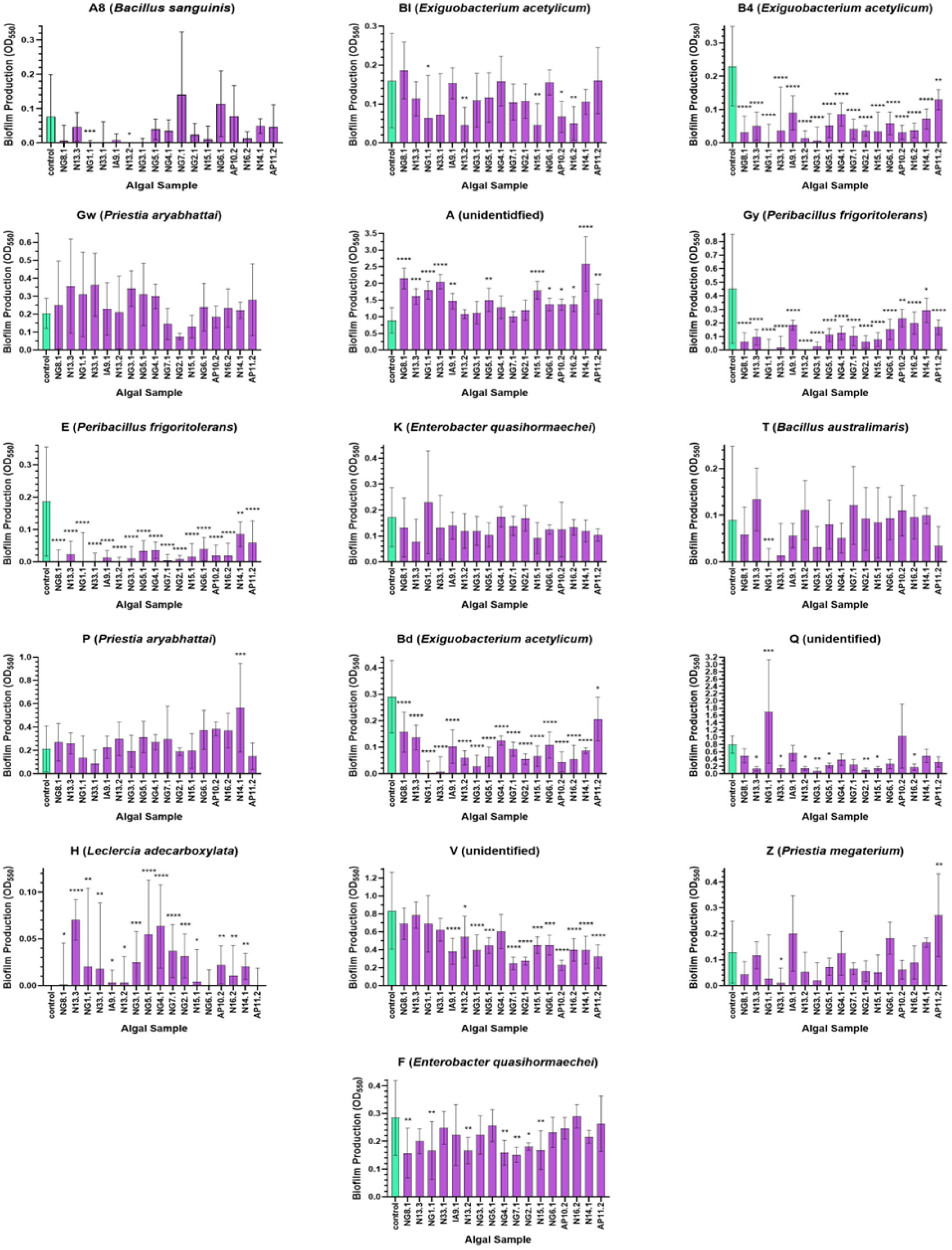
Biofilm production of bacterial isolates. Column graphs show average biofilm production of 16 bacterial isolates over 24 hours, with and without algal supplementation, measured as optical density at 550 nm (OD₅₅₀). Error bars represent ± one standard deviation (n = 3). Data were analysed using a one-way ANOVA followed by Dunnett’s post-hoc test, with statistically significant differences relative to the unsupplemented control indicated by asterisks (* = *p* ≤ 0.05, ** = *p* ≤ 0.01, *** = *p* ≤ 0.001, **** = *p* ≤ 0.0001).

**Supplementary Figure 3.**
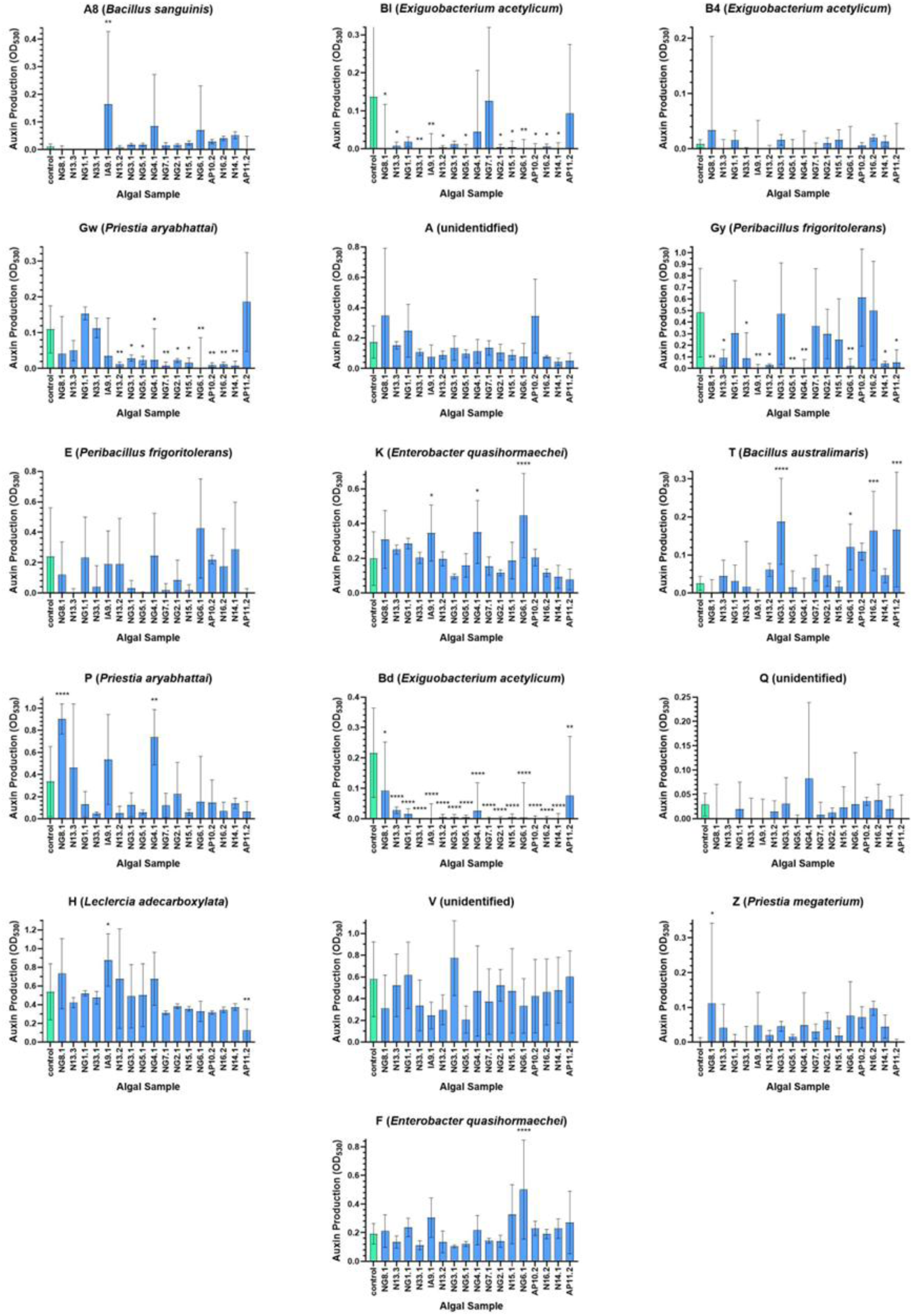
Auxin production of bacterial isolates. Column graphs show auxin production of 16 bacterial isolates over 48 hours, with and without algal supplementation, measured as optical density at 530 nm (OD_530_). Error bars represent ±1 standard deviation (n = 3). Data were analysed using a one-way ANOVA followed by Dunnett’s post-hoc test, with statistically significant differences relative to the unsupplemented control indicated by asterisks (* = *p* ≤ 0.05, ** = *p* ≤ 0.01, *** = *p* ≤ 0.001, **** = *p* ≤ 0.0001).

**Supplementary Figure 4.**
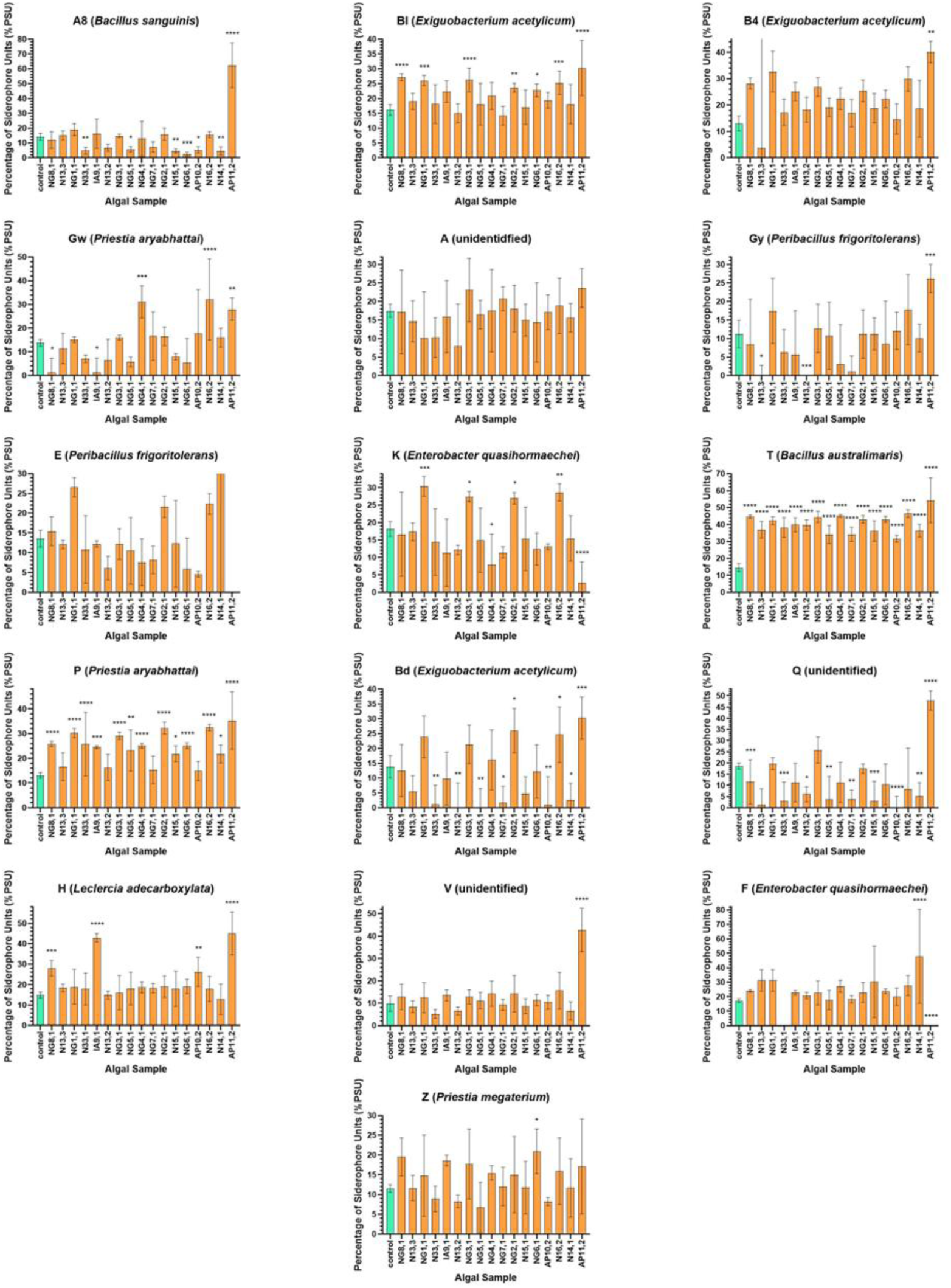
Siderophore production of bacterial isolates. Column graphs show siderophore production of 16 bacterial isolates over 16 hours, with and without algal supplementation, measured as percentage siderophore units (PSU) at 630nm (OD_630_). Error bars represent ± one standard deviation (n = 3). Data were analysed using a one-way ANOVA followed by Dunnett’s post-hoc test, with statistically significant differences relative to the unsupplemented control indicated by asterisks (* = *p* ≤ 0.05, ** = *p* ≤ 0.01, *** = *p* ≤ 0.001, **** = *p* ≤ 0.0001).

**Figure 5.**
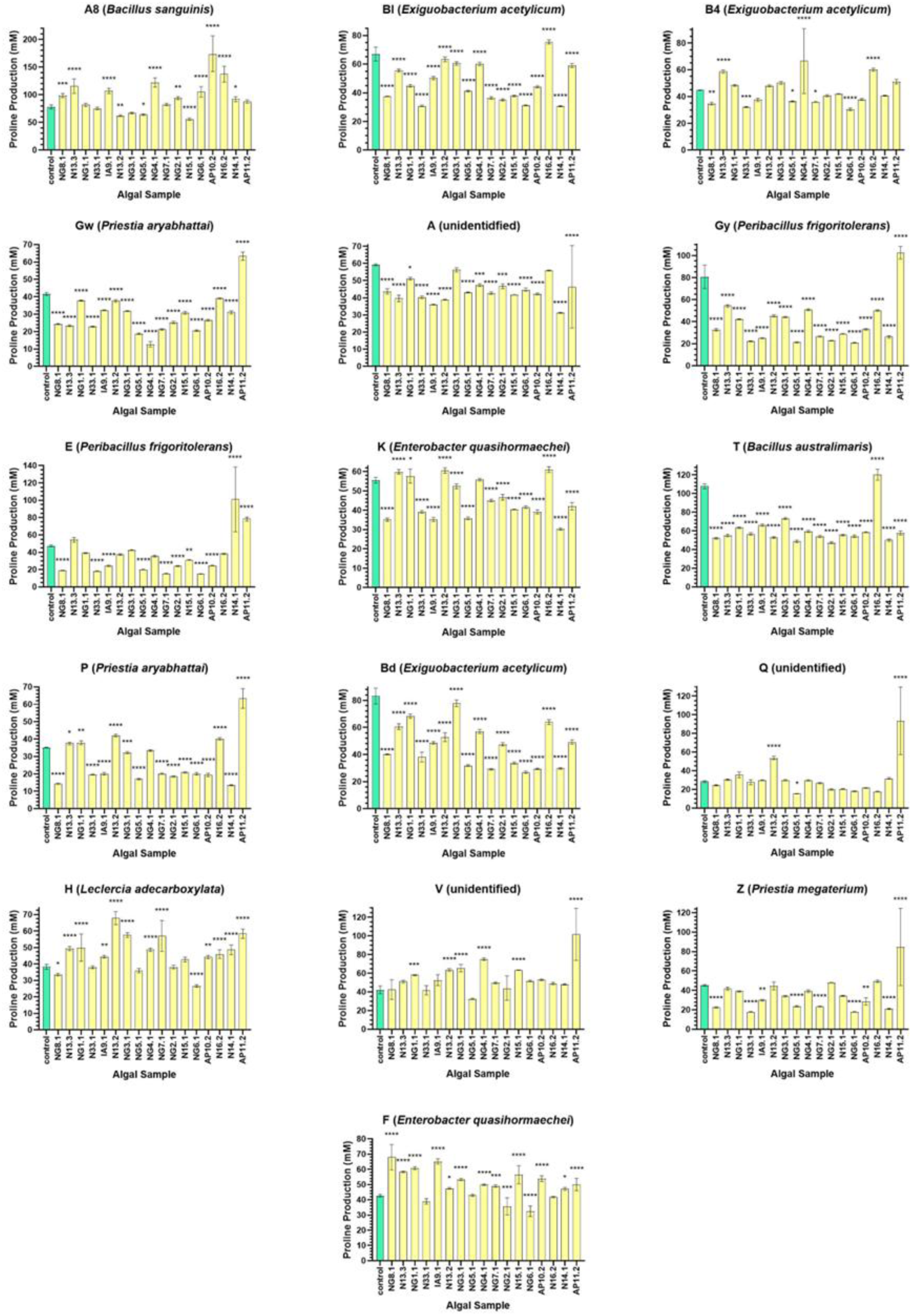
Proline production of bacterial isolates. Column graphs show proline production (mM) of 16 bacterial isolates over two days, with and without algal supplementation, measured at optical density at 520 nm (OD_520_). Error bars represent ± one standard deviation (n = 3). Data were analysed using a one-way ANOVA followed by Dunnett’s post-hoc test, with statistically significant differences relative to the unsupplemented control indicated by asterisks (* = *p* ≤ 0.05, ** = *p* ≤ 0.01, *** = *p* ≤ 0.001, **** = *p* ≤ 0.0001).

**Figure 6.**
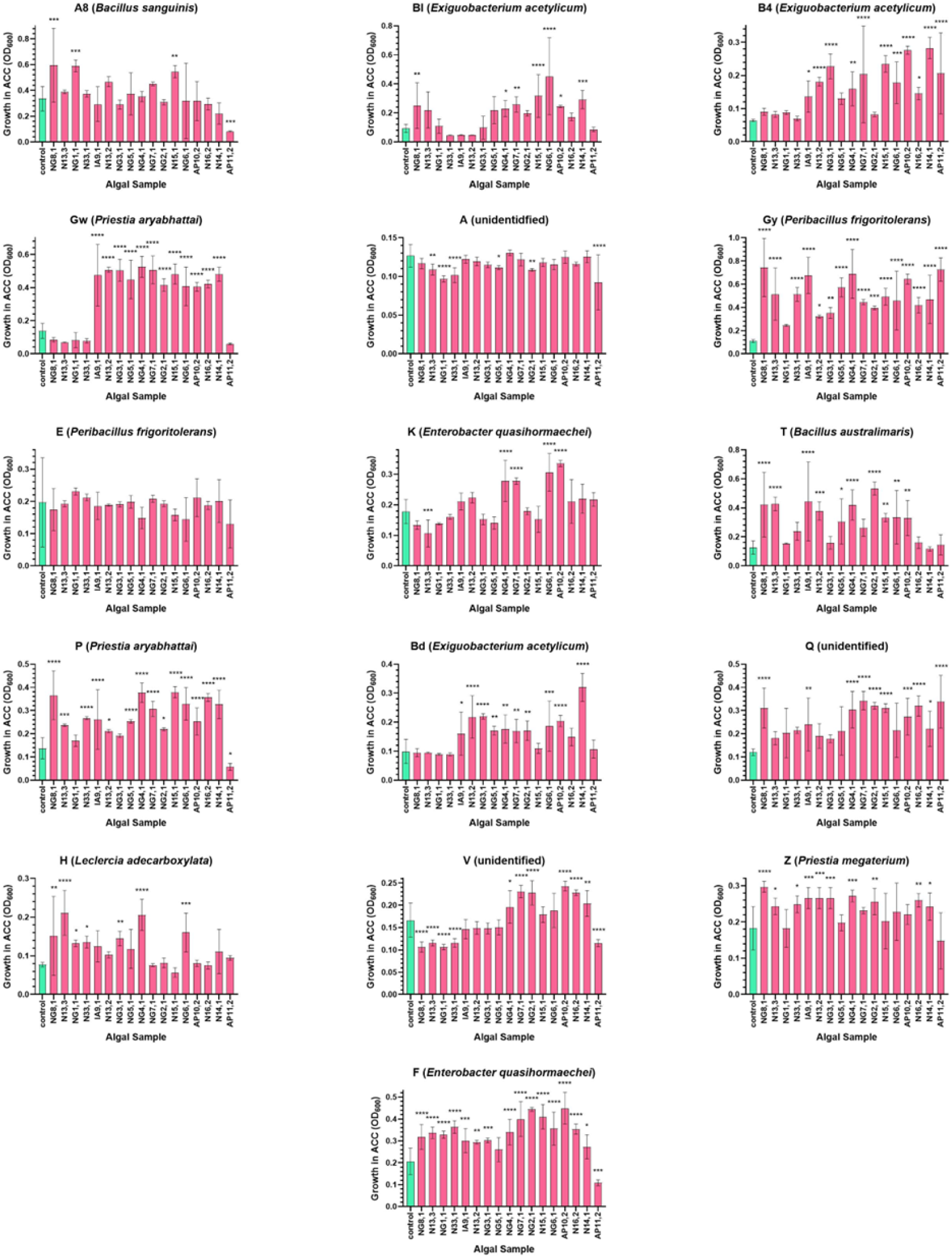
ACC consumption (indirect deaminase activity determination) of bacterial isolates. Column graphs show ACC deaminase activity of 16 bacterial isolates over 3 days, with and without algal supplementation, inferred from growth at 600 nm (OD₆₀₀) with ACC as the sole carbon source. Error bars represent ± one standard deviation (n = 3). Data were analysed using a one-way ANOVA followed by Dunnett’s post-hoc test, with statistically significant differences relative to the unsupplemented control indicated by asterisks (* = *p* ≤ 0.05, ** = *p* ≤ 0.01, *** = *p* ≤ 0.001, **** = *p* ≤ 0.0001).

**Figure 7.**
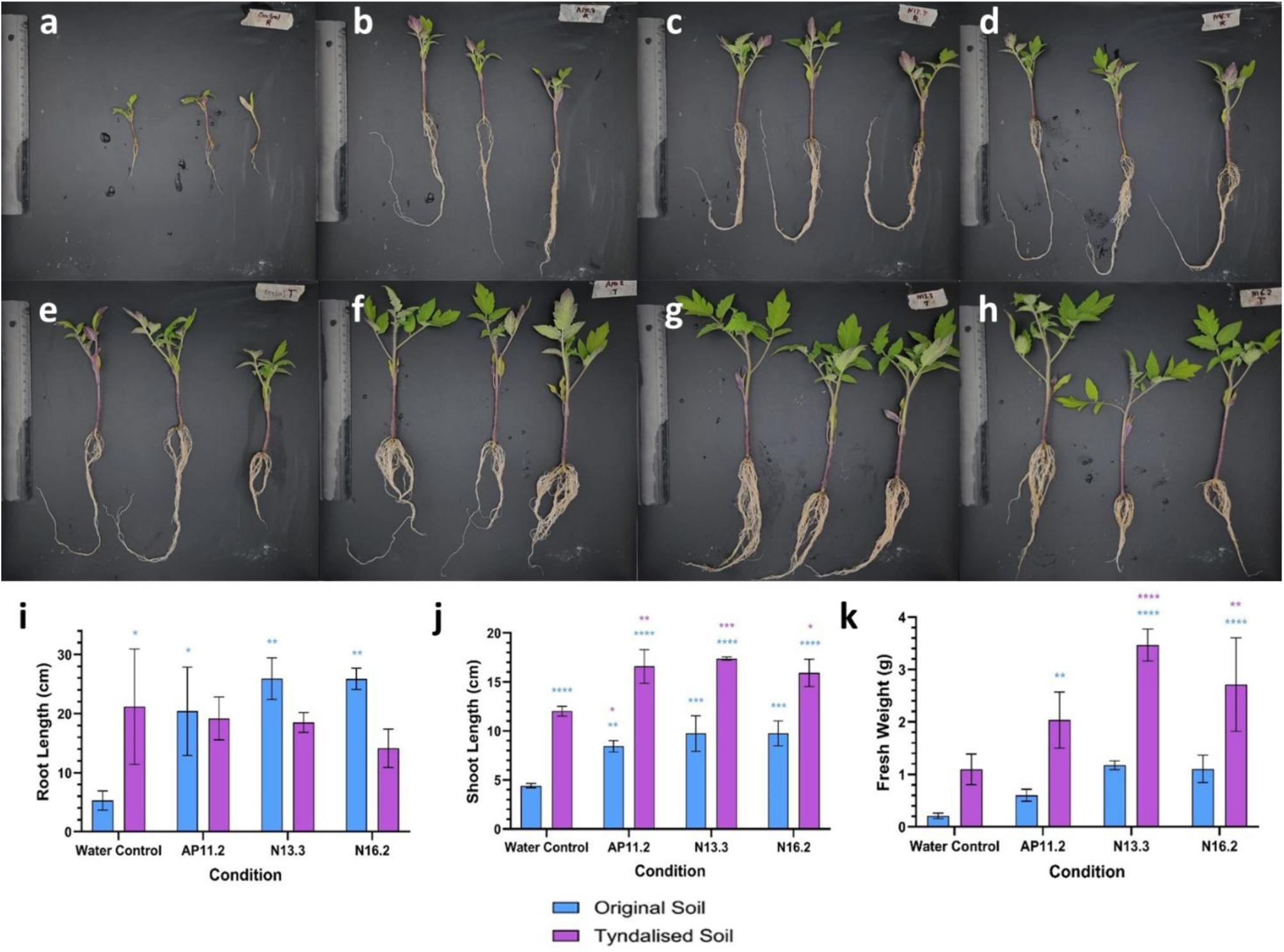
Phenotypic analysis of tomato seedlings under algal application in tyndallized and original soils. Representative triplicates of plants grown in original and tyndallized tomato field soil under different treatments: **(a, e)** untreated control; **(b, f)** AP11.2 extract; **(c, g)** N13.3 extract; and **(d, h)** N16.2 extract. Column graphs show mean **(i)** root length (cm), **(j)** shoot length (cm), and **(k)** dry weight (g) of early-stage tomato plants (n = 3). Statistical analysis was performed using two-way ANOVA to assess the effects of soil type, algal treatment, and their interaction, followed by Tukey’s multiple comparisons test. Blue asterisks indicate significant differences relative to the original soil water control, while purple asterisks indicate significant differences relative to the tyndallized soil water control.

**Supplementary Table 1.** Fold changes in plant growth parameters under algal supplementation relative to water controls. Approximate fold changes in root length (cm), shoot length (cm), and fresh weight (g) were calculated for each algal treatment (AP11.2, N13.3, and N16.2) relative to the corresponding soil-specific water control (original or tyndallized soil).

|  | Root Length |  | Shoot Length |  | Fresh weight |  |
| --- | --- | --- | --- | --- | --- | --- |
|  | Original | Tyndallized | Original | Tyndallized | Original | Tyndallized |
| **AP11.2** | 3.85 | 0.91 | 1.92 | 1.38 | 2.89 | 1.86 |
| **N13.3** | 4.89 | 0.87 | 2.21 | 1.45 | 5.64 | 3.16 |
| **N16.2** | 4.88 | 0.67 | 2.21 | 1.33 | 5.30 | 2.48 |

## ANNEX

**Table A1.**
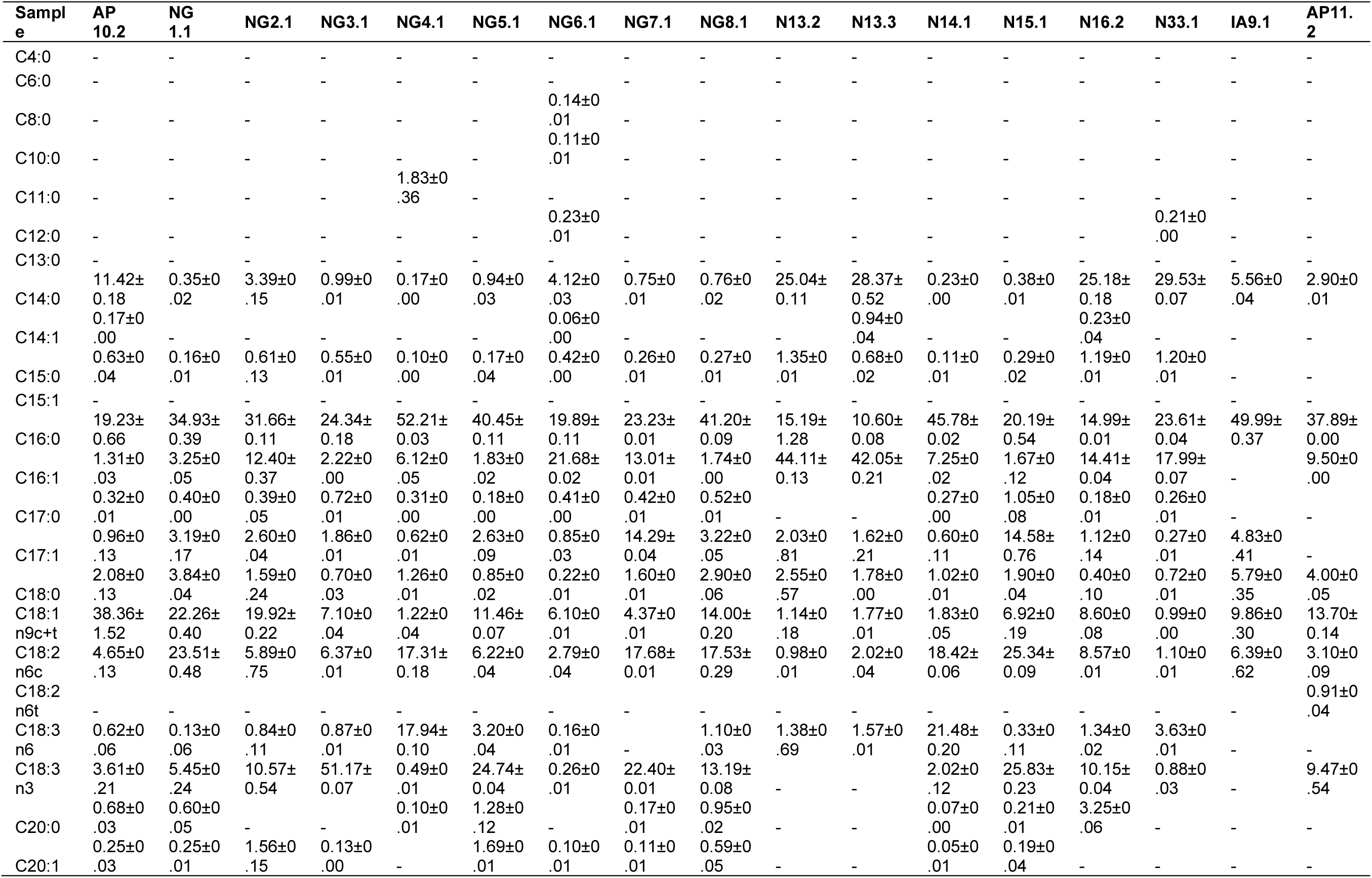
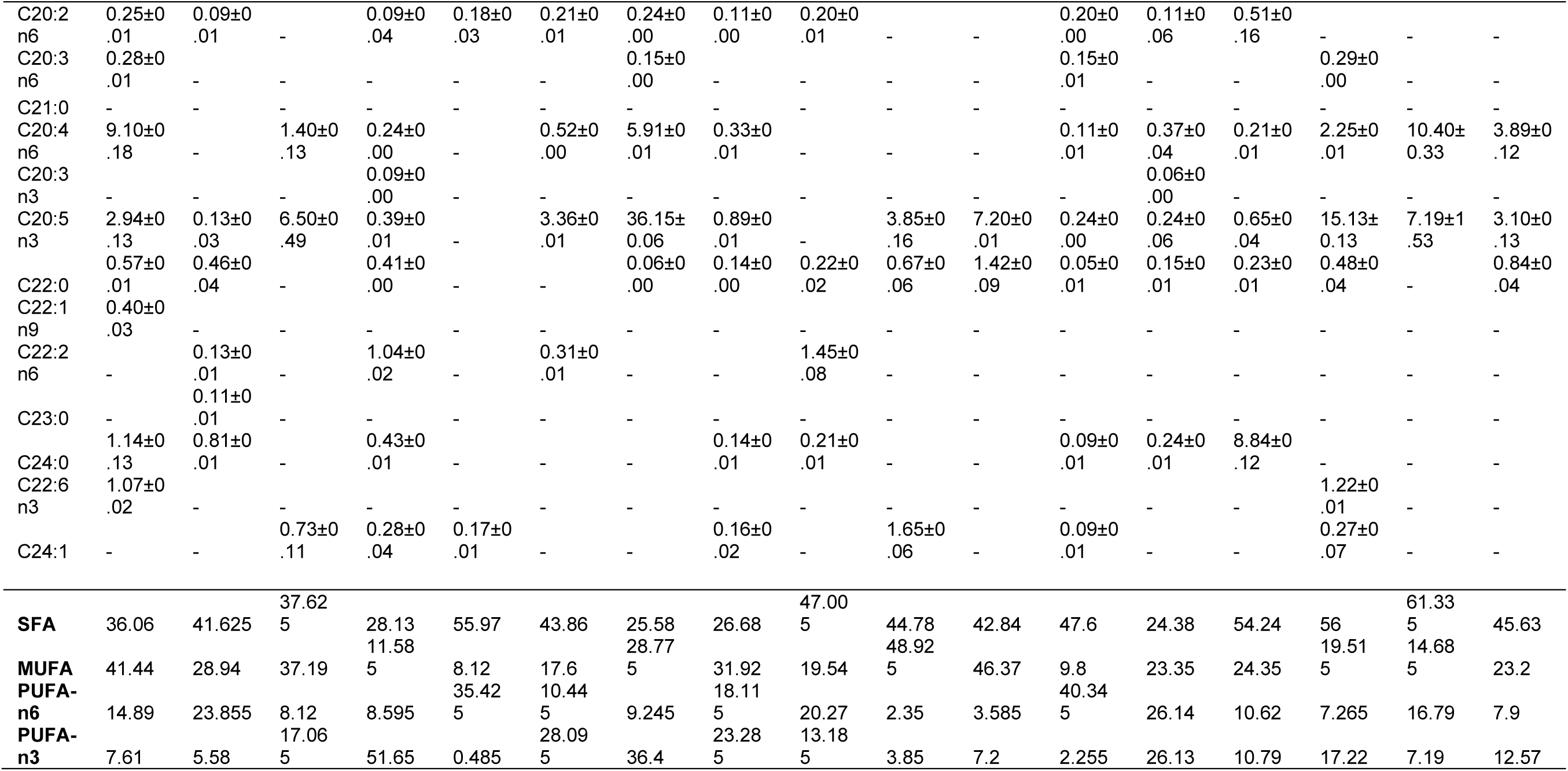
Fatty acid composition determined by GC-FID.

**Table A2.** Volatile compounds isolated by HS-SPME and analyzed by GC-MS.

| Compound | Rt | RI | AP10<br>.2 | NG1.<br>1 | NG2.<br>1 | NG3.<br>1 | NG4.<br>1 | NG5.<br>1 | NG6.<br>1 | NG7.<br>1 | NG8.<br>1 | N13.<br>2 | N13.<br>3 | N14.<br>1 | N15.<br>1 | N16.<br>2 | N33.<br>1 | IA9.<br>1 | AP11<br>.2 |
| --- | --- | --- | --- | --- | --- | --- | --- | --- | --- | --- | --- | --- | --- | --- | --- | --- | --- | --- | --- |
| **ALCOHOLS** |  |  |  |  |  |  |  |  |  |  |  |  |  |  |  |  |  |  |  |
| Ethanol | 1.523 | <700 | 0.46 | 0.22 | 0.26 | 0.64 | 0.75 | 0.33 | 0.34 | 0.79 | 0.37 | 0.44 | 0.57 | - | 1.93 | 0.43 | 0.63 | 2.29 | 0.29 |
| Pentan-1-ol | 2.509 | 769 | 0.09 | 3.09 | 1.67 | - | 0.22 | 1.08 | 1.14 | - | - | 0.07 | - | 0.37 | 0.3 | - | - | - | - |
| Hex-1-en-3-ol | 2.614 | 781 | - | - | - | - | - | - | - | - | - | 1.57 | 2.75 | - | - | - | 1.87 | - | - |
| Hexan-1-ol | 3.762 | 874 | - | - | - | 0.08 | 3.91 | - | - | 0.09 | 0.25 | 0.99 | 0.52 | 3.16 | - | - | - | 0.06 | - |
| (Z)-Octa-1,5-dien-3-ol | 5.97 | 977 | - | - | - | - | - | - | - | - | - | 2.12 | 3.41 | 0.28 | - | 11.21 | 5.43 | - | 0.12 |
| Oct-1-en-3-ol | 6.102 | 982 | 9.56 | 3.48 | - | 3.29 | 0.12 | 4.68 | 4.93 | 3.1 | 4.22 | - | 2.63 | 0.81 | - | 2.61 | 5.22 | 12.29 | 11.25 |
| 2-Ethylhexan-1-ol | 7.595 | 1033 | - | - | - | - | - | - | - | - | - | - | - | 3.47 | 1.28 | - | - | 5.99 | - |
| 2,6-Dimethylcyclohexanol | 10.388 | 1110 | 3.3 | 0.78 | 17.28 | 18.18 | 14.79 | 15.5 | 16.35 | 12.13 | 12.38 | 4.77 | 8.52 | 22.61 | 17.91 | 10.53 | 12 | 3.93 | 2.9 |
| **CARBOXYLIC ACIDS** |  |  |  |  |  |  |  |  |  |  |  |  |  |  |  |  |  |  |  |
| Acetic acid | 1.66 | <700 | 0.16 | 0.11 | 0.82 | 0.14 | 0.07 | 0.03 | 0.03 | 0.31 | 0.2 | - | - | - | 0.31 | - | - | - | - |
| Butanoic acid | 2.588 | 778 | - | - | - | - | - | - | - | 2.41 | 0.53 | 0.21 | - | - | 1.48 | - | - | - | - |
| 3-Methylbutanoic acid | 3.181 | 834 | 0.28 | 1.5 | - | - | 0.21 | - | - | - | 0.67 | 0.4 | 0.23 | - | 1.41 | 0.66 | - | - | 0.19 |
| 2-Methylbutanoic acid | 3.323 | 845 | 0.25 | 0.31 | - | - | 0.05 | - | - | 1.42 | 0.25 | - | - | - | 0.85 | - | - | - | 0.26 |
| Pentanoic acid | 3.827 | 878 | - | 0.64 | - | - | - | - | - | 0.81 | - | - | - | - | 0.43 | 0.29 | - | - | - |
| 4-Methylpentanoic acid | 5.232 | 949 | - | - | - | - | - | - | - | 0.37 | - | - | - | - | - | - | - | - | - |
| 4-Methylpentanoic acid | 5.469 | 958 | - | - | - | - | 0.1 | - | - | 0.46 | - | - | - | - | - | 0.18 | - | - | 0.08 |
| Hexanoic acid | 6.206 | 985 | 8.24 | 22.78 | - | 1.66 | 1.65 | - | - | 6.81 | - | - | - | - | 4.23 | - | - | - | 3.2 |
| Heptanoic acid | 9.334 | 1083 | 3.29 | 3.1 | - | - | - | - | - | 0.73 | - | 2.74 | 2.53 | - | - | - | - | - | 3.5 |
| Octanoic acid | 13.086 | 1181 | 1.93 | 0.83 | - | - | - | - | - | 1.15 | 1.15 | 0.71 | 1.24 | - | - | 0.69 | - | - | 2.9 |
| Nonanoic acid | 17.137 | 1278 | 2.37 | 0.79 | - | 0.31 | - | - | - | 0.33 | 0.79 | 0.33 | - | - | 0.17 | - | - | 1.59 | 2.8 |
| Decanoic acid | 21.122 | 1374 | 0.54 | - | - | - | - | - | - | 0.1 | 0.08 | - | - | - | - | 0.12 | 0.48 | - | 0.22 |
| Dodecanoic acid | 28.925 | 1568 | 0.22 | - | - | - | - | - | - | 0.06 | - | - | - | 1.27 | - | 0.32 | - | 1 | 0.15 |
| Tetradecanoic acid | 36.28<br>4 | 176<br>9 |  |  |  |  |  |  |  |  |  | 0.4 | 0.13 | - | - | 3.15 | 0.51 | - |  |
| **HYDROCARBONS** |  |  |  |  |  |  |  |  |  |  |  |  |  |  |  |  |  |  |  |
| Hexane | 1.727 | <<br>700 | 0.24 | - | - | - | - | - | - | - | - |  |  |  |  |  |  |  | 0.15 |
| p-Xylene | 3.806 | 877 | 0.27 | - | - | 0.51 | - | - | - | - | - | - | - | - | - | - | 0.53 | - | 0.22 |
| Tetradecane | 22.28<br>7 | 140<br>0 | - | - | 0.15 | - | - | - | - | - | - | - | - | - | 0.21 | - | - | - | 0.03 |
| Pentadec-1-ene | 26.00<br>7 | 149<br>2 | 0.1 | - | - | - | - | 0.09 | 0.1 | - | - |  |  |  |  |  |  |  | 0.1 |
| Pentadecane | 26.31<br>4 | 150<br>0 | 0.85 | - | - | 0.84 | 0.73 | - | - | 0.21 | - | - | - | 1.11 | 0.21 | 0.22 | - | - | 0.4 |
| Hexadecane | 30.17<br>4 | 160<br>0 | - | - | - | 0.87 | 1.06 | - | - | 0.14 | - | - | - | - | 0.17 | - | - | 0.62 | 0.51 |
| Heptadec-8-ene | 33.07<br>1 | 167<br>9 | 0.18 | - | - | 2.39 | 0.46 | 0.09 | 0.1 | 0.93 | 1.94 | - | - | 0.55 | 0.94 | - | - | - | 0.22 |
| Heptadecane | 33.86 | 170<br>0 | 0.26 | - | 0.46 | 12.0<br>2 | 27.3<br>3 | 0.16 | 0.16 | 0.95 | 0.32 | 0.24 | - | 27.3<br>8 | 1.87 | - | - | - | 0.18 |
| **ALDEHYDES** |  |  |  |  |  |  |  |  |  |  |  |  |  |  |  |  |  |  |  |
| 3-Methylbutanal | 1.911 | <<br>700 | 0.03 | 0.2 | - | 0.41 | 0.11 | 0.14 | 0.15 | 0.43 | 0.41 | 0.3 | 0.8 | - | 0.56 | 0.52 | 0.51 | 0.11 | 0.13 |
| Pentanal | 2.069 | <<br>700 | 0.24 | 1.34 | 0.53 | 0.84 | 0.15 | 0.23 | 0.25 | 0.51 | 0.46 | 0.21 | 0.34 | - | 1.18 | 0.48 | 0.2 | 0.59 | 0.33 |
| (E)-Pent-2-enal | 2.438 | 761 | 0.09 | 0.11 | - | 0.26 | 0.06 | 0.13 | 0.14 | 0.17 | 0.2 | 0.09 | - | - | - | 0.14 | - | - | 0.08 |
| Hexanal | 2.822 | 801 | 2.13 | 11.1<br>2 | 1.36 | 0.93 | 1.16 | 1.89 | 1.99 | 3.29 | 1.02 | 0.71 | 1.06 | 0.31 | 4.1 | 1.44 | 0.95 | 4.37 | 2.8 |
| 2-Methylpent-2-enal | 3.223 | 837 | - | - | - | 0.14 | - | 0.21 | 0.22 | - | - | - | 0.37 | - | - | - | - | - |  |
| (E)-Hex-2-enal | 3.496 | 857 | 0.1 | 0.22 | 0.33 | 0.77 | 0.1 | 0.46 | 0.48 | 0.28 | 0.46 | 0.44 | 0.52 | - | 0.3 | 0.54 | 0.42 | 0.43 | 0.1 |
| (Z)-Hept-4-enal | 4.26 | 901 | - | - | 0.46 | 0.27 | - | 0.31 | 0.33 | - | 0.66 | - | - | - | 0.17 | 0.3 | 0.17 | - |  |
| Heptanal | 4.292 | 902 | 0.5 | 1.73 | 0.37 | 0.28 | 0.22 | 0.3 | 0.32 | 0.77 | 0.5 | 1.76 | 1.72 | 0.14 | 0.78 | 0.47 | 0.51 | 1.32 | 0.6 |
| (E)-Hept-2-enal | 5.513 | 960 | 0.41 | 0.27 | - | - | 0.07 | - | - | 0.25 | - | 0.34 | 0.23 | - | 0.38 | 0.18 | 0.15 | 1.62 | 0.39 |
| Octanal | 6.762 | 100<br>4 | 0.62 | 2.57 | 0.7 | 0.5 | 0.12 | 0.55 | 0.58 | 0.56 | 0.63 | 0.64 | 0.43 | - | - | - | - | 0.89 | 0.71 |
| (E,E)-Hepta-2,4-dienal | 7.015 | 101<br>3 | 0.47 | 0.29 | 1.32 | 0.82 | 0.12 | 0.64 | 0.67 | 0.39 | 0.94 | 0.4 | 0.33 | - | 0.62 | 0.69 | 0.32 | 0.54 | 0.28 |
| (E)-Oct-2-enal | 8.526 | 106<br>2 | 1.27 | 0.97 | - | - | 0.31 | - | - | 1.35 | - | 0.87 | - | - | - | - | - | 3.57 | 1.78 |
| Nonanal | 10.18<br>6 | 110<br>5 | 1.52 | 4.06 | 1.25 | 0.68 | - | 0.16 | 0.16 | - | - | - | - | - | - | - | - | - | 1.2 |
| (E,Z)-Nona-2,6-dienal | 12.07<br>3 | 115<br>7 | 0.29 | 0.03 | 0.39 | 0.13 | 0.06 | 0.5 | 0.52 | 0.1 | 0.71 | 1.14 | 0.62 | - | - | 0.67 | 0.83 | 1.48 | 0.24 |
| (E)-Non-2-enal | 12.32<br>7 | 116<br>3 | 0.74 | 0.07 | - | 0.06 | 0.23 | 0.21 | 0.22 | 0.26 | 0.42 | 0.66 | 0.37 | - | - | - | - | 0.86 | 0.87 |
| Decanal | 14.20<br>3 | 120<br>6 | 0.12 | 0.34 | 0.82 | 0.14 | 0.26 | 0.14 | 0.15 | 0.15 | 0.16 | - | - | - | 0.19 | - | - | 0.36 | 0.1 |
| (E,E)-Nona-2,4-dienal | 14.54<br>3 | 121<br>5 | 0.4 | 0.83 | - | - | 0.09 | - | - | - | - | - | - | - | 0.05 | - | - | 0.26 | 0.32 |
| (E)-Dec-2-enal | 16.52<br>6 | 126<br>4 | 0.89 | 0.63 | - | - | - | - | - | 0.05 | 0.55 | 0.8 | - | - | 0.38 | 0.63 | - | - | 0.98 |
| (E,Z)-Deca-2,4-dienal | 17.87<br>5 | 129<br>4 | 0.83 | 0.11 | - | - | 0.36 | - | - | 0.19 | - | - | - | - | - | - | - | - | 0.54 |
| (E,E)-Deca-2,4-dienal | 18.83 | 131<br>8 | 1.53 | 1.3 | - | - | 0.41 | 0.09 | 0.1 | 0.05 | 0.37 | - | 2.21 | - | 0.42 | 0.34 | - | 1.28 | 1.2 |
| Dodecanal | 22.66<br>1 | 140<br>9 |  |  |  |  |  |  |  |  |  | 0.36 | - | - | - | - | - | - |  |
| **ESTERS** |  |  |  |  |  |  |  |  |  |  |  |  |  |  |  |  |  |  |  |
| Ethyl but-2-enoate | 3.347 | 847 | - | - | - | - | 0.11 | - | - | 2.12 | - |  |  |  |  |  |  |  |  |
| Ethyl 3-hydroxybutyrate | 4.985 | 938 | - | - | - | - | - | - | - | 0.58 | - |  |  |  |  |  |  |  |  |
| Ethyl hexanoate | 6.634 | 999 | 0.88 | 0.07 | - | 0.46 | 0.22 | 0.17 | 0.18 | 0.75 | 0.13 | - | 0.73 | - | 0.52 | - | - | - | 0.58 |
| Ethyl octanoate | 13.85<br>9 | 119<br>8 | 0.18 | 0.03 | - | 0.01 | 0.02 | 0.22 | 0.23 | 0.2 | 0.38 | 0.14 | 0.4 | - | - | - | - | - | 0.11 |
| Methyl tetradecanoate | 34.74<br>2 | 172<br>5 | 1.38 | - | - | 0.45 | - | - | - | 0.49 | 0.53 | 0.2 | - | - | - | - | - | - | 1.28 |
| Ethyl tetradecanoate | 37.21<br>9 | 179<br>5 |  |  |  |  |  |  |  |  |  | 1.2 | 1.16 | - | - | 1 | 0.47 | - |  |
| **KETONES** |  |  |  |  |  |  |  |  |  |  |  |  |  |  |  |  |  |  |  |
| Heptan-2-one | 4.087 | 892 | 0.1 | 0.34 | - | 0.09 | 0.22 | - | - | 0.69 | 0.16 | - | 0.2 | 0.41 | 0.4 | - | 0.21 | 0.49 |  |
| 6-Methylheptan-2-one | 5.467 | 958 | 0.06 | - | - | 0.37 | 0.1 | 0.25 | 0.26 | 0.46 | 0.2 | 0.21 | 0.53 | 0.25 | - | 0.18 | 0.3 | - | 0.11 |
| Octan-2-one | 6.414 | 992 | 0.01 | 0.03 | - | 0.06 | 0.11 | 0.03 | 0.03 | 1.95 | 0.51 | 0.97 | 1.18 | 0.58 | - | - | - | - | 0.02 |
| 2,2,6-Trimethylcyclohexanone | 7.805 | 104<br>0 | 0.37 | - | 0.54 | 2.86 | 0.44 | 1.19 | 1.25 | 1.67 | 2.18 | 0.37 | 1.75 | 2.07 | 1.34 | 1.21 | 1.63 | - | 0.15 |
| (E)-Oct-3-en-2-one | 7.891 | 104<br>3 | 0.79 | 3.59 | - | - | 0.28 | - | - | 1.15 | - | 0.11 | - | - | - | - | - | - | 0.82 |
| (E,E)-Octa-3,5-dien-2-one | 8.978 | 107<br>4 | 2.92 | 2.03 | 0.93 | 3.7 | 1.1 | 11.4<br>1 | 12.0<br>4 | 5 | 3.48 | 4.3 | 6.34 | 0.68 | 5.86 | 8.09 | 8.28 | 4.31 | 3.2 |
| (E,Z)-Octa-3,5-dien-2-one | 9.79 | 109<br>5 | 3.75 | 10.3<br>4 | 10.1<br>6 | 10.4<br>1 | 0.2 | 12.8<br>1 | 13.5<br>1 | 14.1<br>9 | 5.33 | 5.1 | 4.86 | 1.21 | 10.0<br>5 | 9.23 | 7.52 | 1.86 | 2.9 |
| (E)-Non-3-en-2-one | 11.53<br>5 | 114<br>3 | 0.16 | 0.18 | - | 0.18 | 0.06 | 0.22 | 0.23 | 0.23 | 0.22 | 0.4 | 0.56 | - | - | - | - | - |  |
| 3,5-Nonadien-2-one isomer* | 12.63<br>5 | 141<br>7 |  |  |  |  |  |  |  |  |  | 9.13 | 8.78 | - | - | - | 7.25 | - | 0.22 |
| 3,5-Nonadien-2-one isomer* | 13.67<br>4 | 119<br>4 |  |  |  |  |  |  |  |  |  | 8.22 | 6.84 | - | - | - | 7.69 | - |  |
| Benzenoid derivatives |  |  |  |  |  |  |  |  |  |  |  |  |  |  |  |  |  |  |  |
| Benzaldehyde | 5.691 | 967 | 0.09 | 1.09 | 0.79 | 1.77 | 0.54 | 0.76 | 0.8 | 3.06 | 1.02 | 1.47 | 1.72 | 2.03 | 4.21 | 2.96 | 2.34 | 2.52 | 0.11 |
| Benzeneacetaldehyde | 8.12 | 105<br>0 | 0.33 | - | - | - | 0.43 | - | - | 0.42 | 0.98 | 0.2 | 0.19 | - | - | 0.91 | 0.17 | 1.16 | 0.18 |
| 1-Phenylethanone<br>(Acetophenone) | 8.871 | 107<br>1 | 0.18 | - | - | 1.53 | 0.39 | 2.1 | 2.21 | 0.32 | 4.42 | 0.4 | - | - | - | - | - | 5.2 | 0.05 |
| **APOCAROTENOIDS AND OTHER OXYGENATED TERPENOIDS** |  |  |  |  |  |  |  |  |  |  |  |  |  |  |  |  |  |  |  |
| 6-Methylhept-5-en-2-one | 6.307 | 989 | 0.95 | 3.05 | 9.89 | 0.94 | 0.46 | 7.22 | 7.61 | 1.52 | 1.42 | 0.43 | 0.63 | 1.5 | 1.91 | 2.27 | 0.95 | 2.06 | 0.75 |
| Fenchone | 9.654 | 109<br>1 | 0.99 | - | - | - | - | - | - | - | - | 0.53 | 0.78 | 0.3 | 0.75 | 0.28 | 0.75 | 3.71 | 0.8 |
| 6-Methylhepta-3,5-dien-2-one | 10.22<br>7 | 110<br>6 | - | - | - | 0.74 | 0.78 | 0.94 | 0.99 | 1.62 | 1.45 | 0.93 | 1.47 | 0.68 | 2.2 | 1.71 | 1.22 | 1.46 |  |
| 2,6,6-Trimethyl-<br>cyclohex-2-ene-1,4-dione (4-Oxoisophorone) | 11.72<br>6 | 114<br>8 | 0.53 | 0.72 | 0.51 | 1.52 | 0.68 | 0.56 | 0.59 | 1.44 | 1.57 | 1.72 | 1.49 | 0.6 | 1.14 | 0.29 | 0.59 | 0.06 |  |
| Safranal | 13.98 | 120<br>0 | 0.18 | 0.78 | 0.42 | 1.15 | 1.58 | 0.66 | 0.7 | 1.17 | 2.47 | 0.4 | 0.4 | 0.42 | 1.81 | 1.16 | 0.27 | - | 0.2 |
| β-Cyclocitral | 14.83<br>9 | 122<br>3 | 0.75 | 0.94 | 1.47 | 2.21 | 1.75 | 1.88 | 1.98 | 1.09 | 1.09 | 0.31 | 1.15 | 1.74 | 3.63 | 3.85 | 1.16 | 0.44 | 0.91 |
| α-Ionone | 23.42<br>5 | 142<br>9 | 0.39 | 0.16 | 6.61 | 3.1 | 0.51 | 7.79 | 8.22 | 2.89 | 7.29 | - | - | 0.34 | 4.6 | - | - | 7.18 | 0.74 |
| (E)-<br>Geranylacetone | 24.44<br>5 | 145<br>5 | 0.24 | 0.36 | 0.33 | 0.29 | 0.93 | 0.2 | 0.21 | 0.25 | 0.42 | - | - | - | 0.64 | - | - | - |  |
| 7,8-Epoxy-α-ionone | 25.03<br>6 | 146<br>9 | 0.07 | - | 0.43 | 0.27 | - | 0.38 | 0.4 | 0.19 | 0.58 | - | - | - | 0.15 | - | - | 0.6 | 0.17 |
| β-Ionone | 25.76<br>9 | 148<br>7 | 3.33 | 0.72 | 16.8<br>1 | 12.6<br>4 | 22.3<br>5 | 11.6<br>6 | 12.3 | 5.38 | 12.3 | 8.08 | 9.8 | 16.5<br>3 | 9.1 | 6.19 | 7.07 | 5.57 | 3.2 |
| Dihydroactinidiolide | 27.42<br>3 | 152<br>9 | 2.81 | 0.2 | 6.19 | 3.6 | 7.96 | 3.77 | 3.97 | 3.16 | 6.45 | 3.14 | 2.62 | 5.54 | 3.9 | 4.83 | 3.21 | 5.73 | 3.09 |
| Hexahydrofarnesyl<br>acetone | 38.91<br>8 | 184<br>6 | 0.1 | - | 0.29 | 1.91 | 0.12 | 0.2 | 0.21 | 0.38 | 1.34 | 8.18 | 3.09 | 0.17 | 0.15 | 0.52 | 0.47 | - | 0.09 |
| **OTHERS** |  |  |  |  |  |  |  |  |  |  |  |  |  |  |  |  |  |  |  |
| Dimethyl sulfide | 1.603 | <<br>700 | - | - | 1.86 | - | - | - | - | - | - | - | 1.18 | - | - | 0.8 | - | - | 0.05 |
| Diisobutyl<br>phthalate | 39.69 | 186<br>9 | 0.47 | - | - | 0.34 | - | 0.25 | 0.26 | 0.17 | 0.4 | - | 0.47 | - | - | - | 0.18 | 0.43 | 0.32 |

**Table A3.** Spearman correlation coefficients between individual fatty acids and biofilm formation.

| Fatty Acid | A -<br>not<br>identi<br>fied | A8 -<br>Bacill<br>us<br>sang<br>uinis | B4 -<br>Exiguoba<br>cterium<br>acetylicu<br>m | Bd -<br>Exiguoba<br>cterium<br>acetylicu<br>m | Bl -<br>Exiguoba<br>cterium<br>acetylicu<br>m | E -<br>Peribaci<br>llus<br>frigoritol<br>erans | F -<br>Enterobac<br>ter<br>quasihor<br>maechei | Gw -<br>Priesti<br>a<br>aryab<br>hattai | Gy -<br>Peribaci<br>llus<br>frigoritol<br>erans | H -<br>Leclercia<br>adecarb<br>oxylata | K -<br>Enterobac<br>ter<br>quasihor<br>maechei | P -<br>Priesti<br>a<br>aryab<br>hattai | Q -<br>not<br>identi<br>dfied | T -<br>Bacillu<br>s<br>australi<br>maris | V -<br>not<br>identi<br>dfied | Z -<br>Priesti<br>a<br>megat<br>erium |
| --- | --- | --- | --- | --- | --- | --- | --- | --- | --- | --- | --- | --- | --- | --- | --- | --- |
| C10:0 | -0.12 | 0.37 | 0.19 | 0.19 | 0.25 | 0.31 | 0.19 | 0.0 | 0.19 | -0.37 | 0.0 | 0.37 | 0.12 | 0.12 | 0.12 | 0.31 |
| C11:0 | -0.19 | 0.06 | 0.31 | 0.25 | 0.31 | 0.25 | -0.31 | 0.19 | 0.12 | 0.37 | 0.43 | 0.06 | 0.25 | -0.25 | 0.25 | 0.19 |
| C12:0 | 0.11 | 0.08 | 0.06 | -0.15 | 0.02 | -0.05 | 0.31 | 0.3 | -0.11 | -0.29 | 0.08 | -0.01 | -0.07 | -0.2 | 0.31 | -0.05 |
| C14:0 | -0.04 | -0.29 | -0.13 | -0.3 | -0.2 | -0.34 | 0.47 | 0.29 | -0.21 | -0.17 | -0.02 | -0.22 | -0.39 | 0.06 | 0.22 | -0.06 |
| C14:1 | 0.03 | 0.33 | 0.1 | 0.13 | -0.02 | 0.29 | 0.32 | 0.22 | 0.29 | 0.07 | -0.19 | 0.35 | -0.19 | 0.49 | 0.33 | 0.29 |
| C15:0 | -0.18 | -0.41 | **-0.72** | **-0.61** | **-0.6** | **-0.56** | 0.04 | 0.06 | **-0.61** | 0.05 | -0.05 | -0.05 | **-0.74** | 0.28 | 0.38 | **-0.61** |
| C16:0 | 0.22 | -0.01 | 0.42 | 0.26 | **0.56** | 0.21 | -0.13 | 0.04 | 0.27 | 0.04 | 0.34 | -0.09 | **0.62** | **-0.54** | -0.21 | 0.23 |
| C16:1 | -0.28 | 0.19 | -0.1 | -0.15 | -0.3 | -0.11 | 0.1 | 0.1 | -0.19 | 0.04 | -0.04 | 0.18 | -0.39 | 0.49 | 0.23 | 0.01 |
| C17:0 | -0.06 | -0.01 | -0.47 | -0.14 | -0.06 | -0.17 | -0.41 | -0.22 | -0.28 | 0.05 | 0.1 | -0.04 | -0.15 | -0.14 | -0.07 | **-0.51** |
| C17:1 | -0.22 | -0.16 | -0.39 | -0.04 | -0.2 | -0.44 | -0.51 | -0.5 | -0.25 | 0.09 | 0.08 | 0.01 | -0.02 | 0.25 | -0.19 | -0.32 |
| C18:0 | 0.14 | -0.16 | 0.12 | **0.54** | 0.2 | -0.18 | -0.43 | -0.29 | -0.09 | -0.27 | -0.1 | -0.35 | 0.36 | 0.01 | -0.07 | 0.16 |
| C18:1<br>n9c+t | 0.05 | 0.01 | 0.02 | 0.17 | 0.41 | 0.1 | 0.2 | -0.23 | 0.2 | -0.26 | 0.04 | -0.18 | 0.15 | -0.21 | -0.47 | 0.12 |
| C18:2<br>n6c | 0.19 | 0.2 | 0.05 | 0.15 | -0.05 | 0.19 | -0.36 | -0.41 | 0.38 | 0.11 | 0.12 | 0.24 | 0.4 | 0.05 | -0.27 | 0.01 |
| C18:2<br>n6t | 0.12 | 0.19 | 0.43 | 0.43 | 0.37 | 0.37 | 0.37 | 0.12 | 0.25 | -0.43 | -0.31 | -0.37 | 0.19 | -0.31 | -0.31 | 0.43 |
| C18:3<br>n3 | -0.03 | -0.05 | -0.42 | -0.3 | -0.14 | -0.11 | -0.01 | -0.12 | -0.18 | 0.14 | -0.14 | -0.15 | -0.27 | -0.23 | -0.37 | -0.49 |
| C18:3<br>n6 | 0.34 | -0.12 | -0.07 | -0.25 | -0.14 | 0.1 | 0.02 | 0.41 | -0.09 | 0.48 | -0.08 | 0.28 | -0.1 | 0.03 | **0.58** | -0.21 |
| C20:0 | 0.19 | 0.13 | -0.07 | 0.0 | -0.11 | 0.09 | -0.1 | -0.15 | 0.26 | 0.17 | 0.03 | 0.46 | 0.27 | 0.15 | 0.06 | -0.13 |
| C20:1 | -0.01 | 0.08 | -0.39 | -0.14 | 0.05 | -0.1 | -0.21 | -0.27 | -0.24 | 0.14 | -0.09 | 0.05 | -0.16 | 0.0 | -0.22 | -0.39 |
| C20:2<br>n6 | 0.11 | 0.35 | 0.09 | 0.02 | 0.08 | 0.42 | 0.14 | -0.04 | 0.46 | 0.0 | 0.03 | **0.76** | 0.33 | 0.18 | 0.08 | 0.13 |
| C20:3<br>n3 | -0.07 | -0.37 | **-0.51** | -0.33 | -0.3 | -0.1 | -0.03 | -0.01 | -0.33 | 0.01 | -0.35 | -0.32 | -0.46 | -0.29 | -0.1 | -0.46 |
| C20:3<br>n6 | 0.37 | 0.18 | 0.16 | -0.19 | -0.09 | 0.12 | 0.29 | 0.2 | 0.09 | -0.2 | -0.01 | 0.16 | 0.07 | -0.09 | 0.23 | 0.02 |
| C20:4<br>n6 | -0.01 | 0.14 | 0.37 | -0.1 | 0.11 | 0.1 | **0.52** | -0.08 | 0.24 | -0.36 | 0.1 | -0.32 | 0.06 | -0.38 | -0.49 | 0.28 |
| C20:5<br>n3 | -0.14 | 0.08 | 0.14 | -0.15 | -0.07 | -0.14 | 0.38 | 0.16 | -0.12 | -0.09 | -0.03 | -0.17 | -0.3 | 0.11 | 0.04 | 0.15 |
| C22:0 | 0.13 | -0.24 | -0.32 | 0.02 | -0.2 | -0.22 | 0.16 | 0.36 | -0.37 | -0.25 | -0.53 | -0.3 | -0.39 | 0.07 | 0.33 | -0.24 |
| C22:1<br>n9 |  |  |  |  |  |  |  |  |  |  |  |  |  |  |  |  |
| C22:2<br>n6 | 0.1 | -0.34 | -0.42 | -0.04 | 0.37 | -0.19 | -0.02 | 0.37 | -0.33 | 0.03 | -0.17 | -0.02 | -0.01 | -0.35 | 0.15 | -0.45 |
| C22:6<br>n3 | 0.31 | -0.31 | -0.12 | -0.43 | -0.25 | -0.43 | 0.25 | 0.43 | -0.37 | 0.0 | 0.12 | -0.43 | -0.25 | -0.43 | 0.31 | -0.43 |
| C23:0 |  |  |  |  |  |  |  |  |  |  |  |  |  |  |  |  |
| C24:0 | 0.05 | -0.17 | -0.48 | -0.26 | -0.34 | -0.13 | -0.08 | -0.21 | 0.02 | -0.06 | -0.08 | 0.15 | -0.1 | 0.07 | -0.16 | -0.39 |
| C24:1 | **-0.51** | -0.37 | -0.45 | **-0.55** | -0.37 | -0.49 | -0.33 | -0.14 | **-0.56** | 0.27 | 0.38 | -0.21 | -0.44 | -0.06 | -0.14 | -0.47 |
| C8:0 | -0.12 | 0.37 | 0.19 | 0.19 | 0.25 | 0.31 | 0.19 | 0.0 | 0.19 | -0.37 | 0.0 | 0.37 | 0.12 | 0.12 | 0.12 | 0.31 |

**Table A4.**
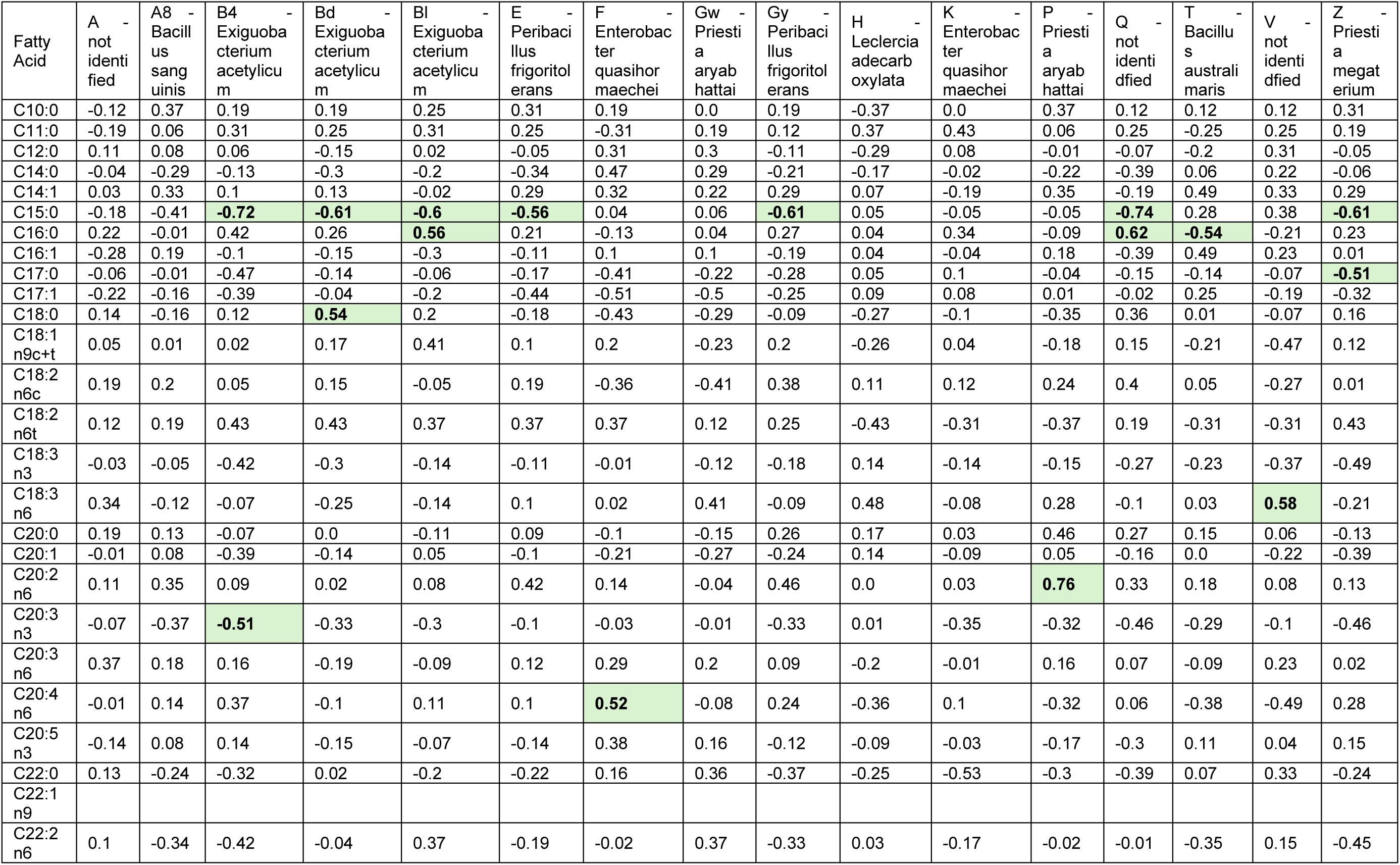
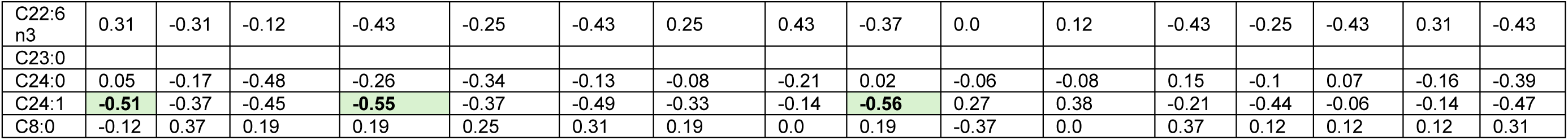
Spearman correlation coefficients between individual fatty acids and biofilm formation.

**Table A5.**
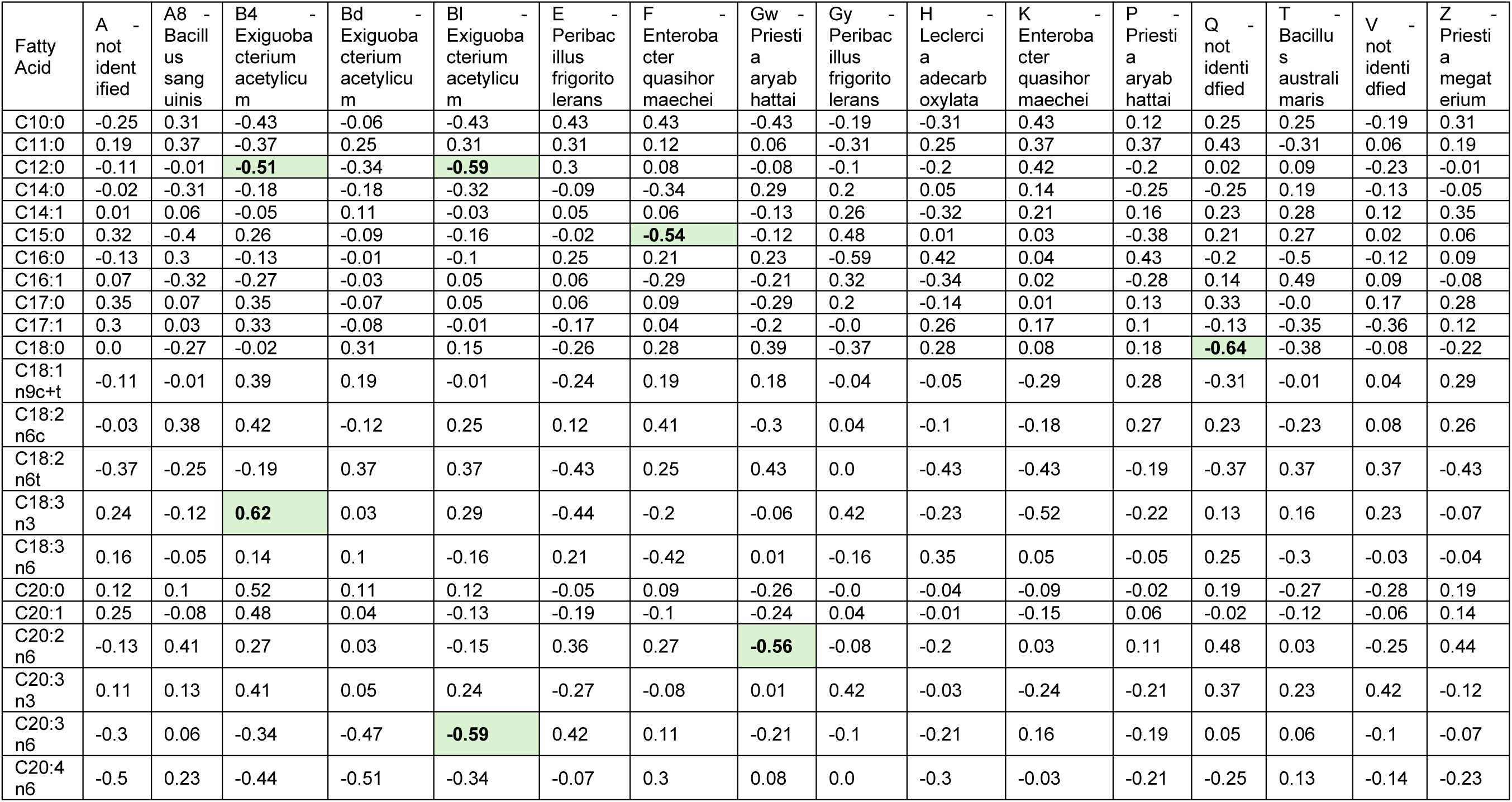
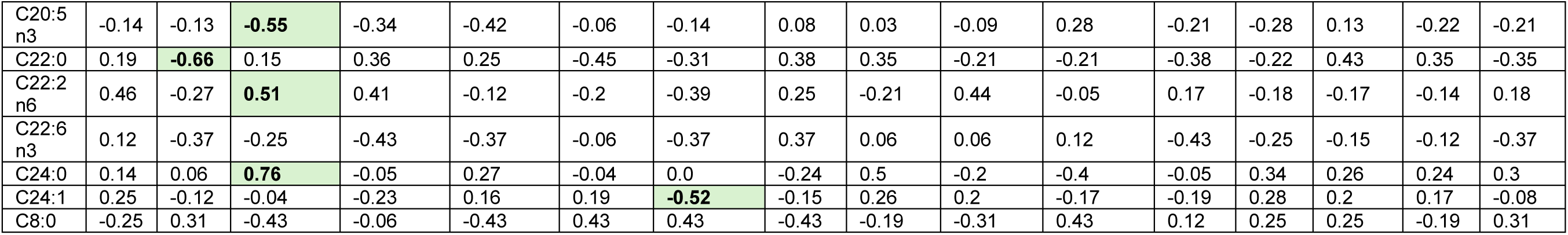
Spearman correlation coefficients between individual fatty acids and auxins production.

**Table A6.**
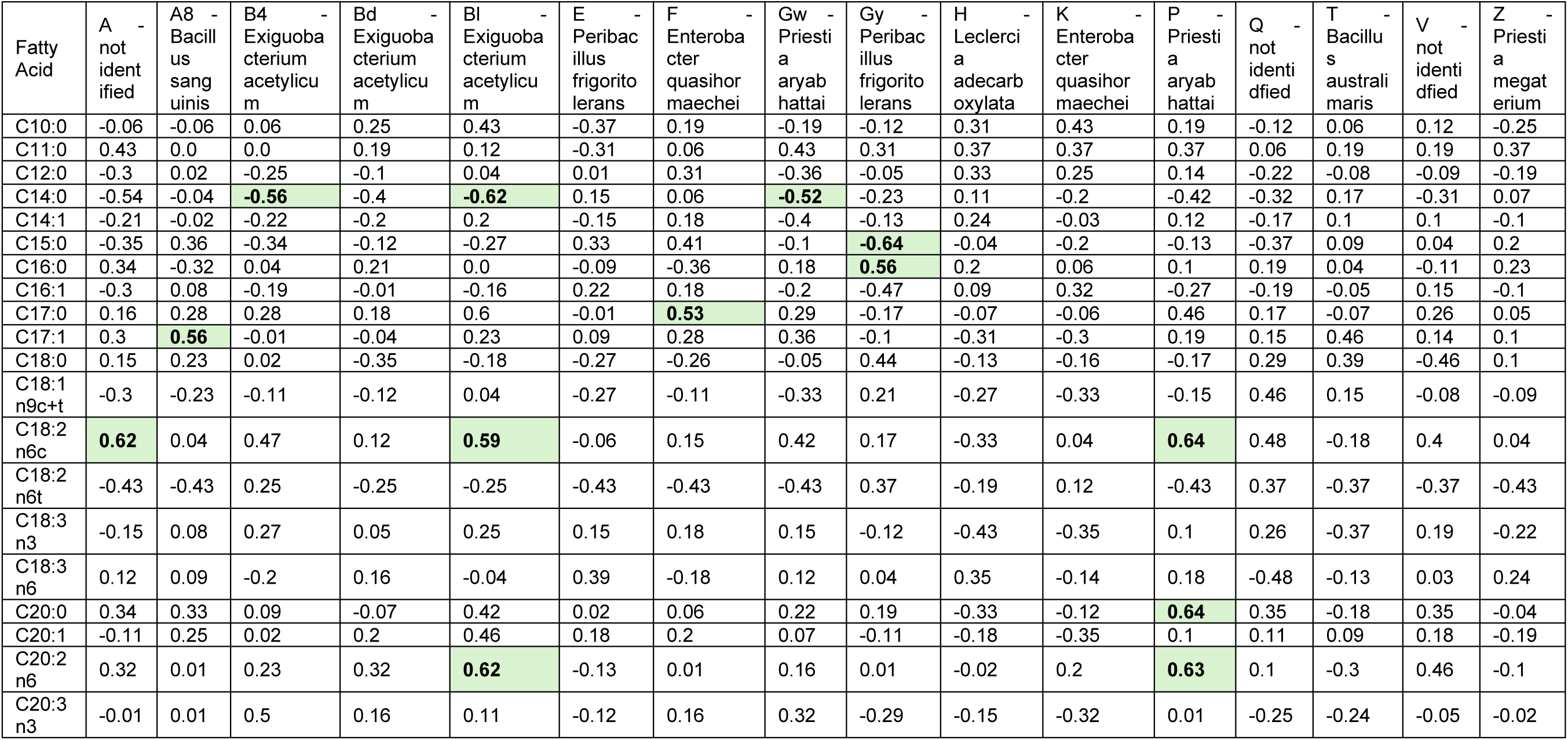
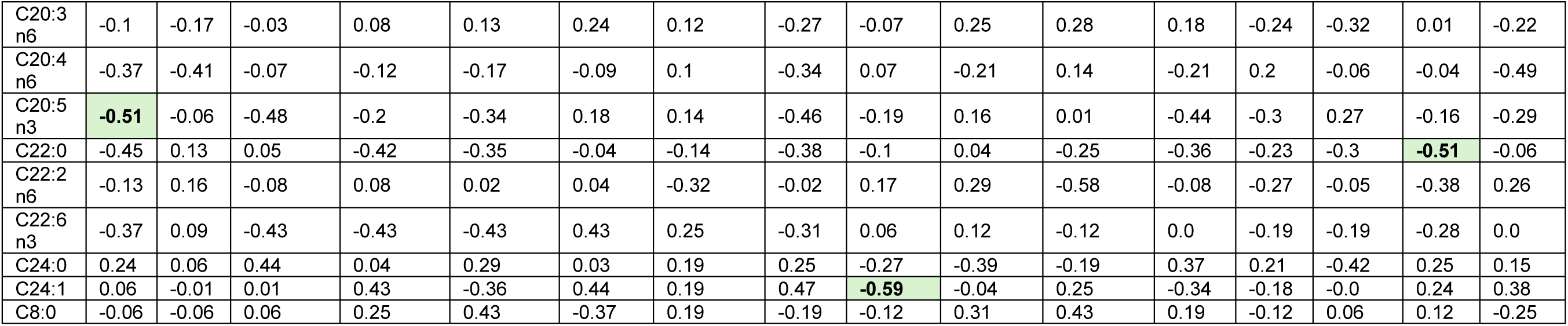
Spearman correlation coefficients between individual fatty acids and ACC deaminase production.

**Table A7.**
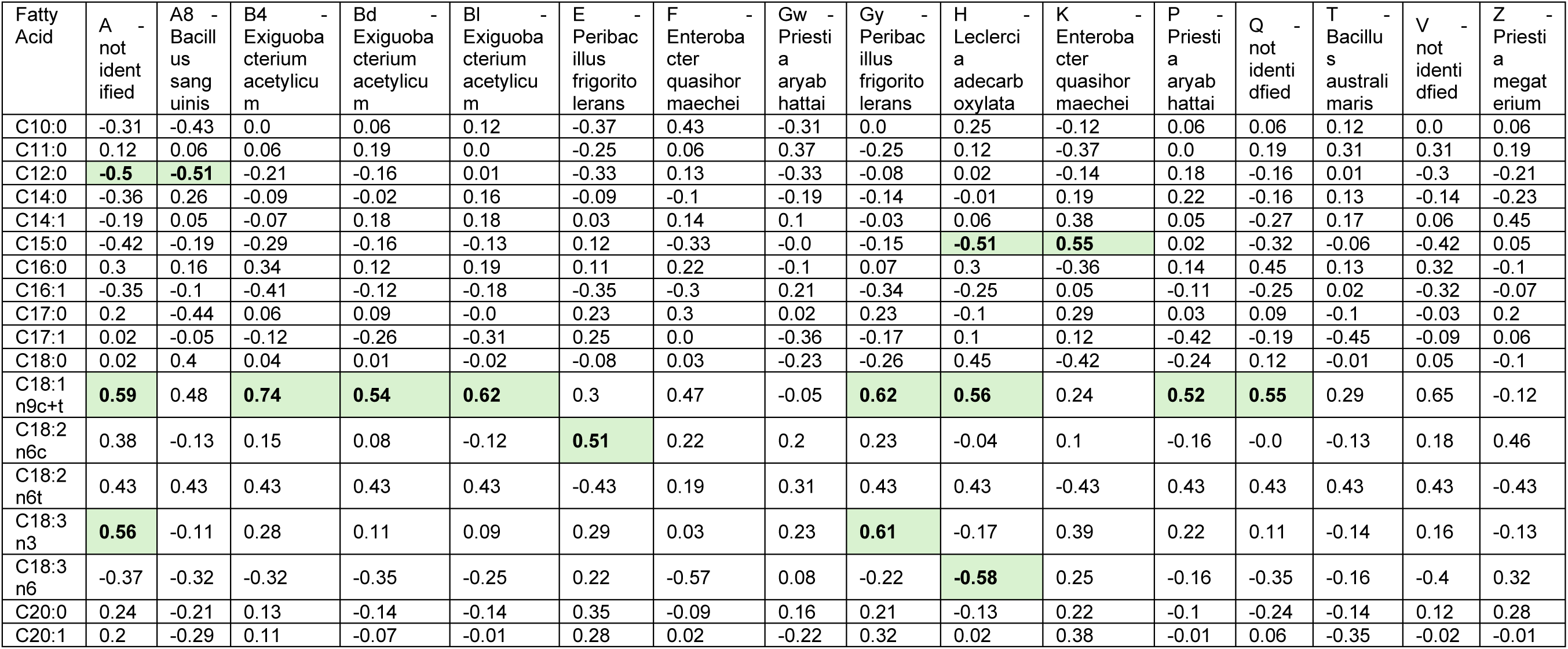
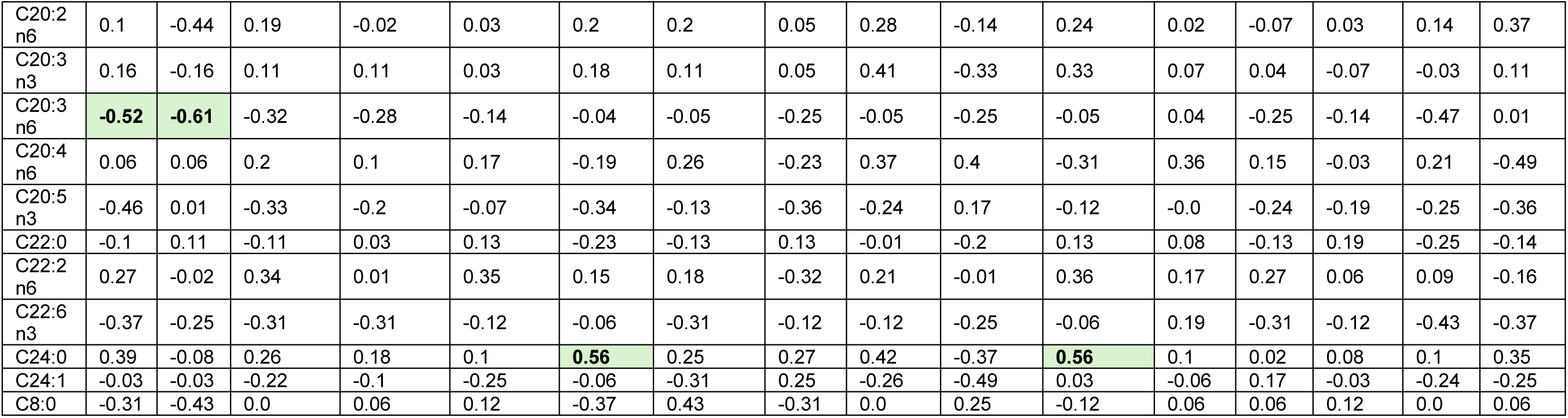
Spearman correlation coefficients between individual fatty acids and siderphore production.

**Table A8.**
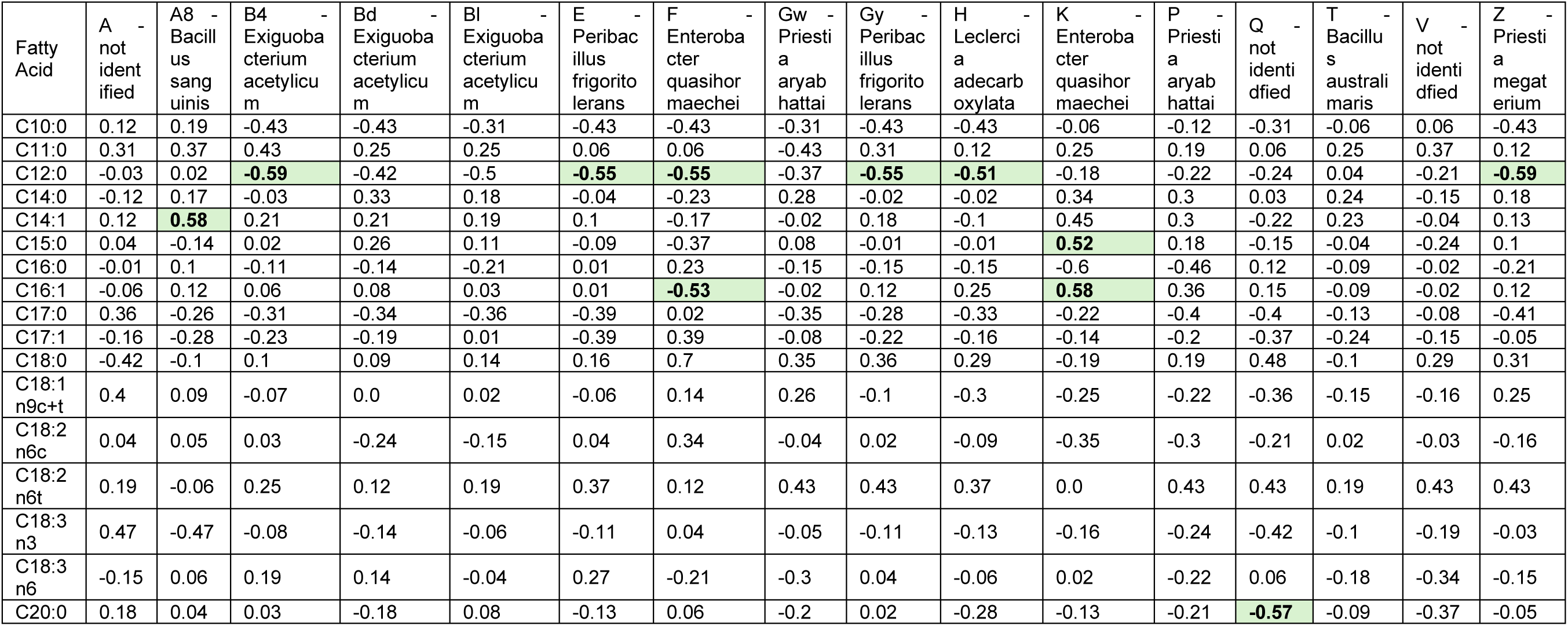
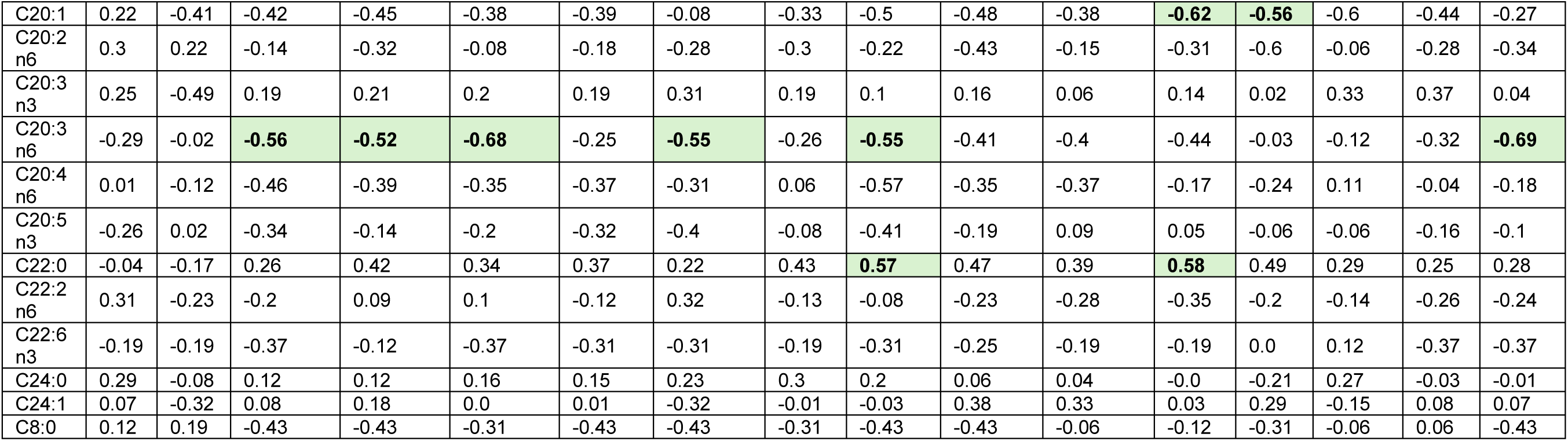
Spearman correlation coefficients between individual fatty acids and proline production.

